# Human tissue-resident CD8 T cells contribute to trophoblast homeostasis in health and during acute inflammation

**DOI:** 10.64898/2026.06.05.730192

**Authors:** Caitlin S. DeJong, Marie Frutoso, Nicole B Potchen, Geervani Daggupati, Andrew J. Konecny, Yin Huang, Manu Setty, Swati Shree, Stephen A McCartney, Martin Prlic

## Abstract

Previous studies have highlighted that some T cell subsets in tissues can provide signals to support tissue cell homeostasis and differentiation. If and how T cell-tissue cell signaling is altered in healthy compared to inflamed tissues is poorly understood. Here, we address if communication between human T cells and tissue cells changes from steady state to an acutely inflamed state in the human placenta. We used single cell analysis strategies to examine invasive cytotrophoblasts (iCTBs) and immune cells isolated from third trimester healthy and acutely inflamed human placentas. We performed cell communication analysis to predict cell-cell communication networks, and found evidence that iCTBs provided signals to support the recruitment of T cells, as well as the formation of tissue-resident memory CD8 T cells (Trm). In exchange, Trm provide signals to support iCTB homeostasis. During acute inflammation, iCTBs and macrophages underwent profound transcriptional changes, while most T cell subsets only underwent limited transcriptional changes. This was not due to T cell exhaustion or tolerance, as T cells were functionally intact. Cell communication analysis and validation at the protein level provide evidence that T cells can maintain their homeostatic support to iCTBs during acute inflammation.

## INTRODUCTION

Injury or infection of tissue cells initiates an immune response that typically includes activation of the innate immune cells followed by the activation of adaptive immune cells(1, 2). While there are canonical responses from tissue cells in response to injury(3) and infection(4), recent human and mouse model studies reported that epithelial cells, endothelial cells and fibroblasts also elicited distinct responses compared to each other and across different organs(4, 5). Similarly, cells of the immune system exert responses in a tissue-dependent manner, which has been shown for macrophages and T cells(6). This includes tissue-resident memory T cells (Trm), which show tissue-associated Trm functional changes that have been referred to as functional adaptation(7–13).

Previous studies have demonstrated that tissue cells and T cells can exchange signals directly. This has been most extensively studied in context of the tumor environment where tumor cells can shape T cell responses via receptor-, cytokine- and metabolite-mediated signals(14). In the context of a tumor, these signals are typically inhibitory and tumor cells have been reported to express high levels of PD-L1, secrete TGFβ, and alter tissue metabolism. Importantly, signals are not exclusively delivered in a unilateral manner from tissue cells to instruct immune cells. In the mouse model, regulatory T cells communicate with hair follicle stem cells to regulate their growth and differentiation in a JAG-1-dependent manner during healthy steady state (SS) and following inflammatory events, highlighting that immune cells can also play a role in maintaining proper organ function(15, 16). This suggests that tissue cells and immune cells have a communication network in place to ensure that they carry out tissue-appropriate functions.

Ideally, the communication network between tissue cells and immune cells would be assessed in the context of SS conditions and compared to a well-defined inflammatory insult. This would allow to establish which networks are maintained and altered in the context of inflammation. We sought to identify a well-controlled cohort of study participants that would enable the study of tissue and immune cells from SS and acutely inflamed tissues. The full-term (third trimester) human placenta affords an unparallelled opportunity to sample tissue and immune cells from both SS and acute inflammation.

Full-term placental tissues are readily available from both SS and those experiencing acute inflammation in the form of intraamniotic inflammation (IAI) providing a defined cohort with acute inflammation.

Here, we interrogate the signals exchanged between invasive cytotrophoblast cells (iCTBs) and T cells within the decidual-placental interface (DPI) within SS and IAI placentas using imaging, Antibody-Sequencing (AbSeq), high parameter flow cytometry, and cell communication analysis tools. In SS conditions, iCTBs provided signals to recruit T cells and promote tissue residence, and T cells provided signals to promote the survival and differentiation of iCTBs. Acute inflammation led to profound changes in iCTBs and antigen presenting cells (APCs) (dendritic cells and macrophages), but T cells showed minimal alterations, with survival and differentiation signals remaining maintained between T cells and iCTBs. Additionally, we demonstrate that T cells from the DPI are responsive to *ex vivo* stimulation, suggesting that they are functionally intact. Overall, our data indicate that tissue and immune cells can maintain mutual support during acute inflammation.

## RESULTS

### T cells and invasive cytotrophoblasts (iCTBs) co-localize within the DPI at steady state

To gain spatial context if invasive cytotrophoblasts (iCTBs) and T cells interact within the 3^rd^ trimester (full term) decidual-placental interface (DPI) we used immunofluorescence to label tissue and immune cells (**Fig 1**). We used cytokeratin-7 (CK7) to visualize the spatial distribution of iCTBs, vimentin (VIM) for mesoderm-derived immune, stem cell, and fibroblast cells, and CD8 for CD8^+^ T cells within the DPI and find their co-localization throughout the tissue. We found that CD8 T cells co-localized next to iCTBs (**Fig 1a,i,b,ii**). We next wanted to gain additional insight to determine if these cells were merely co-localized or potentially interacting. We used transmission electron microscopy (TEM) to visualize subcellular interactions (**Fig 1c,d,iii**). Here we find intertwined projections between iCTBs and cells that appear morphologically consistent with immune cells (**Fig 1d,iii**), suggesting direct cell - cell membrane interactions. Taken together, these microscopy data indicate that iCTBs and immune cells are poised to interact within the full term decidual-placental interface at SS.

**Figure 1:**
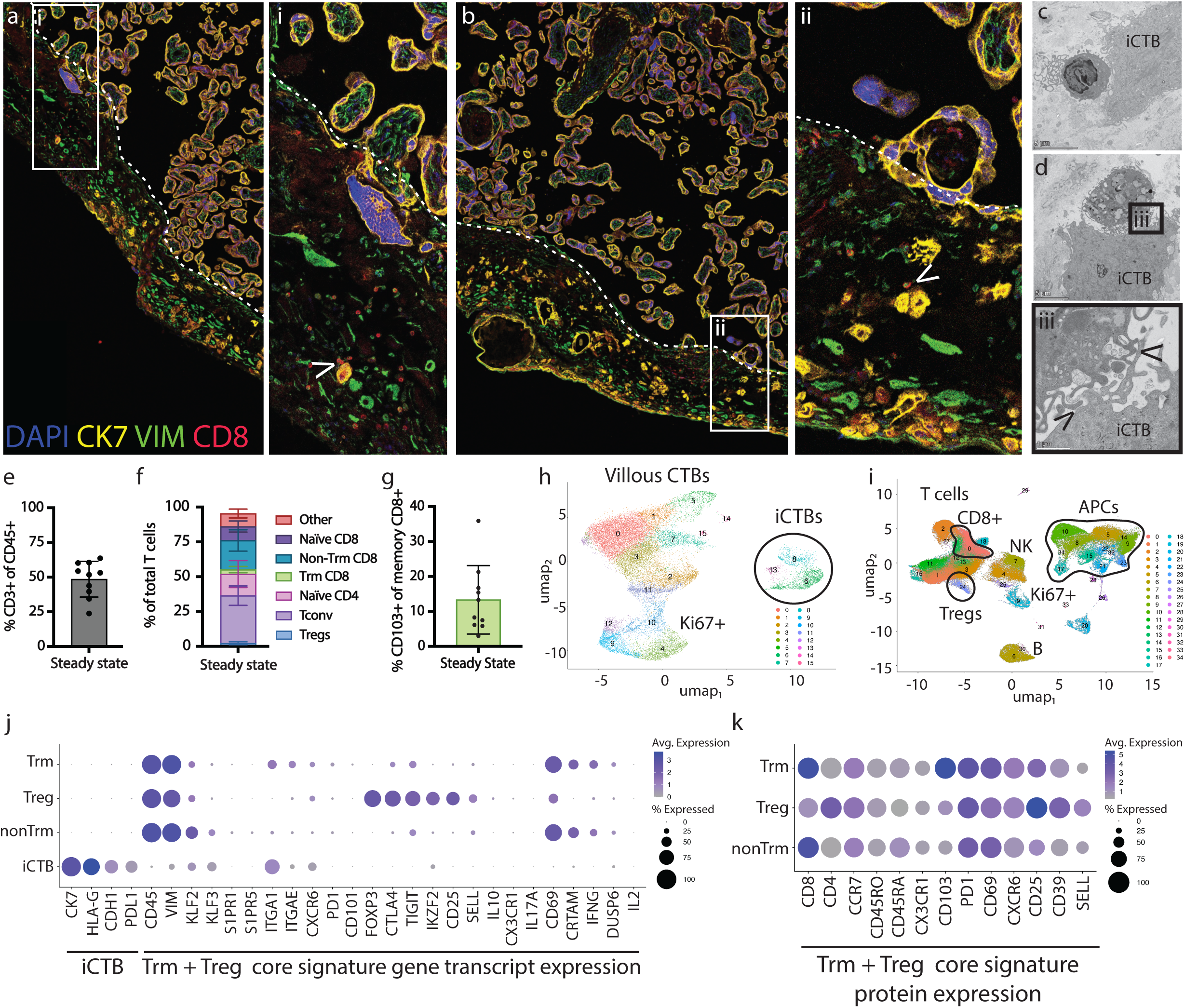
iCTBs co-localize with immune cells and are poised to interact during steady state. **a,b,** Confocal microscopy images from within the decidual-placental interface (DPI). Dashed line indicates where the placental villi and decidua basalis meet. DAPI labels nuclei; Cytokeratin 7 (CK7) specifically labels fetal ectoderm-derived cytotrophoblasts; vimentin (VIM) labels both fetal and maternal mesoderm-derived cells including fibroblasts, stem cells, and immune cells; CD8 specifically labels cytotoxic T lymphocytes including tissue-resident memory T cells. **i,ii,** Zoom-in of the two boxed regions in a and b. Arrowhead indicates colocalization of invasive cytotrophoblasts (iCTBs) with CD8^+^ T cells. **c,d,** Transmission electron microscopy from within the DPI. **c,** Larger cell consistent with an iCTB colocalizing with a cell consistent with an immune cell. **d,** Intertwined projections of an iCTB and a cell consistent with an antigen presenting cell (APC). **iii,** Zoom-in of the boxed region in **d.** Arrowheads indicate points of intercellular membrane contact. **e,f,g,** We used a previously published flow cytometry dataset(17) to assess frequencies within the DPI during steady state of all T cells (**e**), T cell subsets (**f**), and Tissue-resident memory CD8^+^ T cells (**g**), as measured by flow cytometry. Each point represents a study participant (n=10). Error bars are standard deviation (SD). MAIT and gamma delta T cells (Other), CD103^-^ memory CD8 T cells (Non-Trm CD8), CD103^+^ memory CD8 T cells (Trm CD8), conventional CD4^+^ T cells (Tconv), regulatory T cells (Tregs). **h,** Uniform Manifold Approximation and Projection (UMAP) of single-cell integration of cytotrophoblast cells (29,162) from full term placentas at SS (n=4); villous cytotrophoblasts (CTBs), invasive cytotrophoblasts (iCTBs), cycling trophoblasts (Ki67+). **i,** UMAP of single-cell integration of all immune cells (121,857) from full term placentas at SS (n=6), including cells from bloods and DPI. NK cells (NK), CD14^+^ antigen-presenting cells (APCs), B cells (B). **j,** Bubble plot depicting average transcript expression level and frequency of iCTB, Trm, and Treg signature genes. **k,** Bubble plot depicting average protein expression level and frequency of Trm and Treg signature genes.

We next used a flow cytometry dataset from a recent study(17) to determine which T cell subsets populate the DPI during steady state (SS). As previously reported, the immune compartment within the full-term human placenta is comprised of approximately 50% T cells (18, 19) (**Fig 1e**). The T cell compartment consisted of CD4^+^ and CD8^+^ T cell populations, including naïve, memory (non-Trm CD8, Tconv), regulatory T cells (Treg), tissue-resident (Trm CD8), as well as MAIT and γδ T cells (Other) (**Fig 1f**; Supp Fig 1). Furthermore, we found the memory CD8^+^ T cell compartment is comprised of 10-20% CD103^+^ CD8 T cells, which we refer to as Trm (**Fig 1g**).

To interrogate the signal exchange between T cells and iCTBs, we intended to use NicheNet. Briefly, NicheNet uses scRNAseq data to infer cell-cell communication. Importantly, NicheNet goes beyond simply assessing expression of ligands and receptors, and instead also takes expression of receptor target genes into account to infer active signaling events and increase the robustness of these predictions.

To generate the datasets needed for NicheNet, we performed single cell RNA sequencing (scRNAseq) experiments to examine trophoblast cells (n=4 donors; 29,162 cells) and CD45+ immune cells specific to the DPI (n=6 donors; 85,717 cells) from SS (**Fig 1h,i**; Supp Fig 2; Supp Fig 3). Briefly, the CTBs can be divided into two populations: iCTBs (*HLA-G^+^*, *KRT7^+^*, *EGFR^-^*) of the DPI, and villous CTBs (*HLA-G^-^*, *KRT7^+^*, *EGFR^+^*, *KI67^+^*) which compose the placental structure responsible for nutrient, waste, gas exchange, and humoral uptake (20–24). The CD45+ DPI populations contained the following immune cell subsets: T cells (CD3^+^), natural killer (NK) cells (CD56^+^), B cells (CD19^+^), and antigen presenting cells (APCs) including CD14^+^ dendritic cells and macrophages (**Fig 1i**; Supp Fig 3). To ensure our study focused on the T cells in the DPI tissue, cells with signatures present in maternal blood (mBlood) and cord blood (cBlood) were removed from the analysis (Supp Fig 4) to focus on populations exclusively present in the DPI.

As a final quality control step before examining cell – cell communication with NicheNet, we assessed iCTB, Trm, non-Trm, and Treg populations to determine if these cells express expected core transcriptional signatures. We examined the core signature genes (average expression and percent expressed) at both the transcript (**Fig 1j**) and protein level (average expression and percent expressed, using antibody sequencing where possible) (**Fig 1k**). In brief, iCTBs uniquely express cytokeratin 7 (*CK7*), the non-classical MHC Class I protein *HLA-G*, E-caherin (*CDH1*), and programmed cell death ligand 1 (PD-L1/*CD274*). Within the T cell population, Trm will uniquely express *ITGA1* and CD103 (*ITGAE*), important for tissue retention, and *CXCR6* important for homing to a tissue site; their low expression of *KLF2*, *KLF3*, *SELL*, *S1PR1*, and *CX3CR1* supports their avoidance of tissue exit cues (11). *FOXP3*, *CTLA4*, *TIGIT*, and *IKZF2* (*HELIOS*), along with *CD25*, confirmed the Treg population (**Fig 1 j,k**).

### T cells within the DPI are predicted to support homeostasis of iCTBs during steady state

Using NicheNet, we first addressed the types of signals T cells (Trm CD8^+^, non-Trm CD8^+^, and Tregs) were predicted to send to iCTBs. We found most predicted interactions were in support of trophoblast survival and differentiation and not unique for a specific T cell subset, but redundantly provided across T cell subsets (**Fig 2a**). We specifically identified a combination of cytokines and chemokines provided by T cells that were previously demonstrated to support trophoblast survival and differentiation including CSF1, BMP7, XCL1(Lymphotactin), CCL5 (RANTES), and TGFβ1 (25–28).

**Figure 2:**
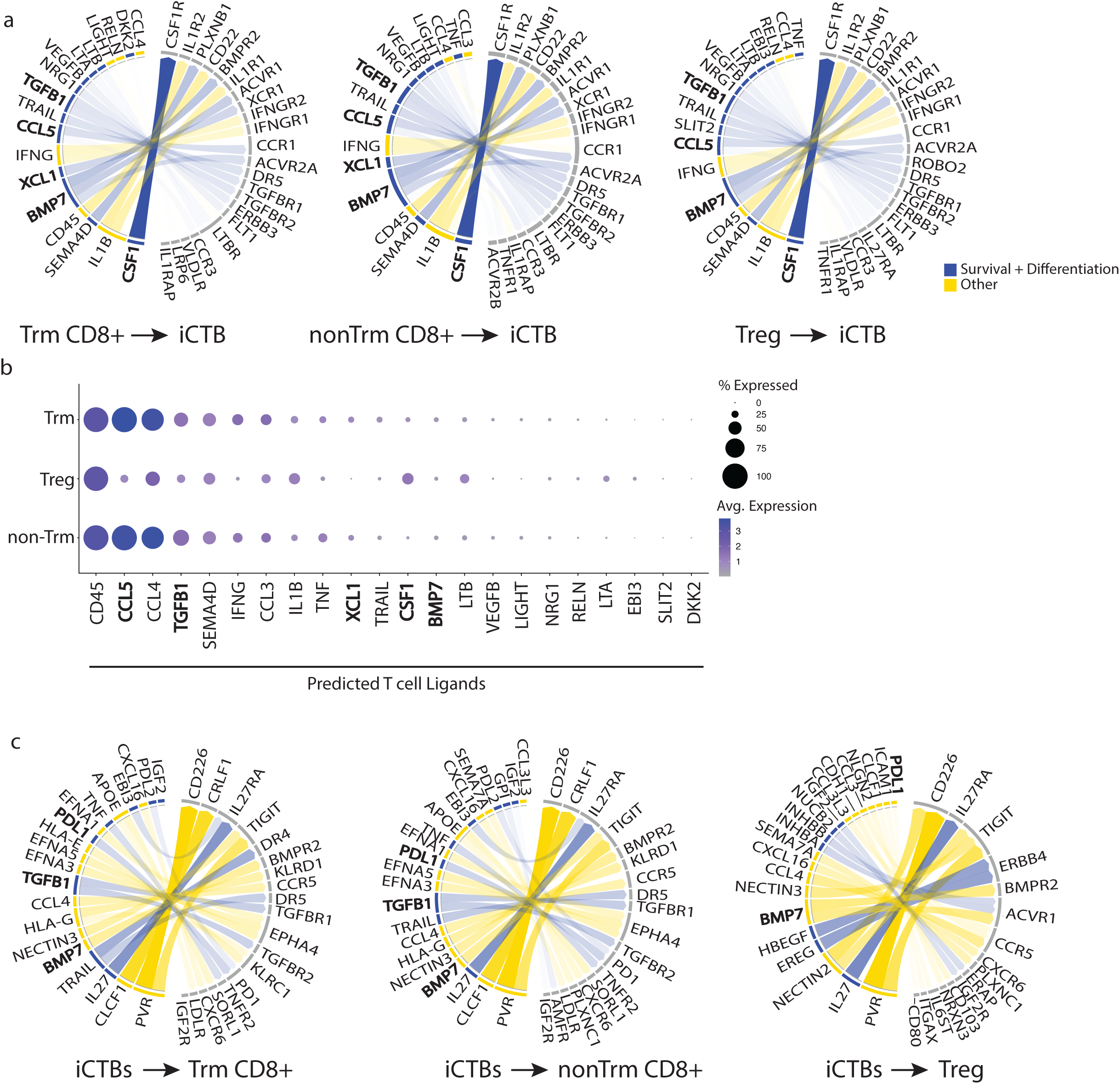
Single cell transcriptome profiling of the DPI during steady state revels potential for a mutual tissue-immune exchange of survival and differentiation factors that is cell-type specific. **a**, Circos plots visualizing the top 25 NicheNet-predicted interactions between T cells and iCTBs during SS. Interactions with a known role in iCTB survival and differentiation are blue, all others are yellow. Ligands are on the left of each plot, receptors of engaged signaling pathways are on the right. Ligands of specific interest are bold. **b,** Bubble plot depicting average transcript expression level and frequency of predicted T cell ligands. **c,** Circos plots visualizing the top 25 NicheNet-predicted interactions between iCTBs and T cells during SS. Interactions with a known role in Trm survival and differentiation are blue, all others are yellow. Ligands are on the left of each plot, receptors of engaged signaling pathways are on the right. Ligands of specific interest are bold.

Next, to determine how specific or redundant the NicheNet-predicted T cell ligands are within the DPI, we visualized the degree of transcript expression across T cell subsets (**Fig 2b**). In doing so, we were able to appreciate that although most ligands are redundantly predicted across the three T cell subsets, the degree of transcript expression associated with promoting iCTB homeostasis varied between Tregs and Trm/nonTrms (e.g. *CCL5*, *TGFβ1*).

We next asked if iCTBs could provide signals to support recruitment and or maintenance of Trm. The predicted cell – cell communication between iCTBs and Trm CD8^+^ during SS was consistent with tissue-supported Trm retention, homeostasis, and fitness; specifically, iCTBs were predicted to provide two key factors for Trm generation and maintenance: TGFβ1 and PD-L1 (29, 30) (**Fig 2c**).

Overall, NicheNet predictions suggested that both T cells and iCTBs express the molecular programs capable of mutually supporting overall survival, differentiation, and homeostasis (blue lines) within the DPI at SS. Lastly, it was previously shown that placental trophoblasts express the adhesion molecule E-cadherin (CDH1), the CD103 ligand, further supporting the potential for iCTB-Trm interactions within the DPI(22).

After determining that iCTBs and T cells exhibit transcriptional programs in support of mutually beneficial survival and fitness, we next wanted to see if this relationship is maintained during inflammation.

### Local and systemic changes associated with intra-amniotic inflammation (IAI)

To examine iCTB – T cell signaling predictions during a state of acute inflammation, we isolated trophoblasts and immune cells from the placentas of subjects with intra-amniotic inflammation (IAI). IAI is diagnosed based on a maternal temperature greater than 38°C with maternal and/or fetal tachycaridia (31).

In an effort to assess the systemic level of inflammation across subjects, we measured levels of IL-6, a key player in the acute inflammatory response (32) which has also been shown to be elevated in the amniotic fluid of subjects with experiencing IAI (33). We examined IL-6 concentrations in mBlood plasma (systemic) and DPI lysates (local). Consistent with acute inflammation, we found that IL-6 levels increased significantly and over 10-fold in IAI (both in DPI tissue lysates and in maternal plasma) compared to SS (**Fig 3a**). Another key indicator of acute inflammation is neutrophil recruitment. We find an increase in the G-CSF, a known neutrophil chemoattractant (34), during IAI when compared to SS, both in DPI lysates and maternal plasma (**Fig 3a**). Taken together, these data indicate that our IAI samples exhibited robust states of acute inflammation in line with their clinical diagnosis. Finally, IL-15 was detected in tissue lysates and maternal plasma, but although IL-15 availability can increase following infections(35), SS levels were overall maintained during IAI milieu (**Fig 3a**).

**Figure 3:**
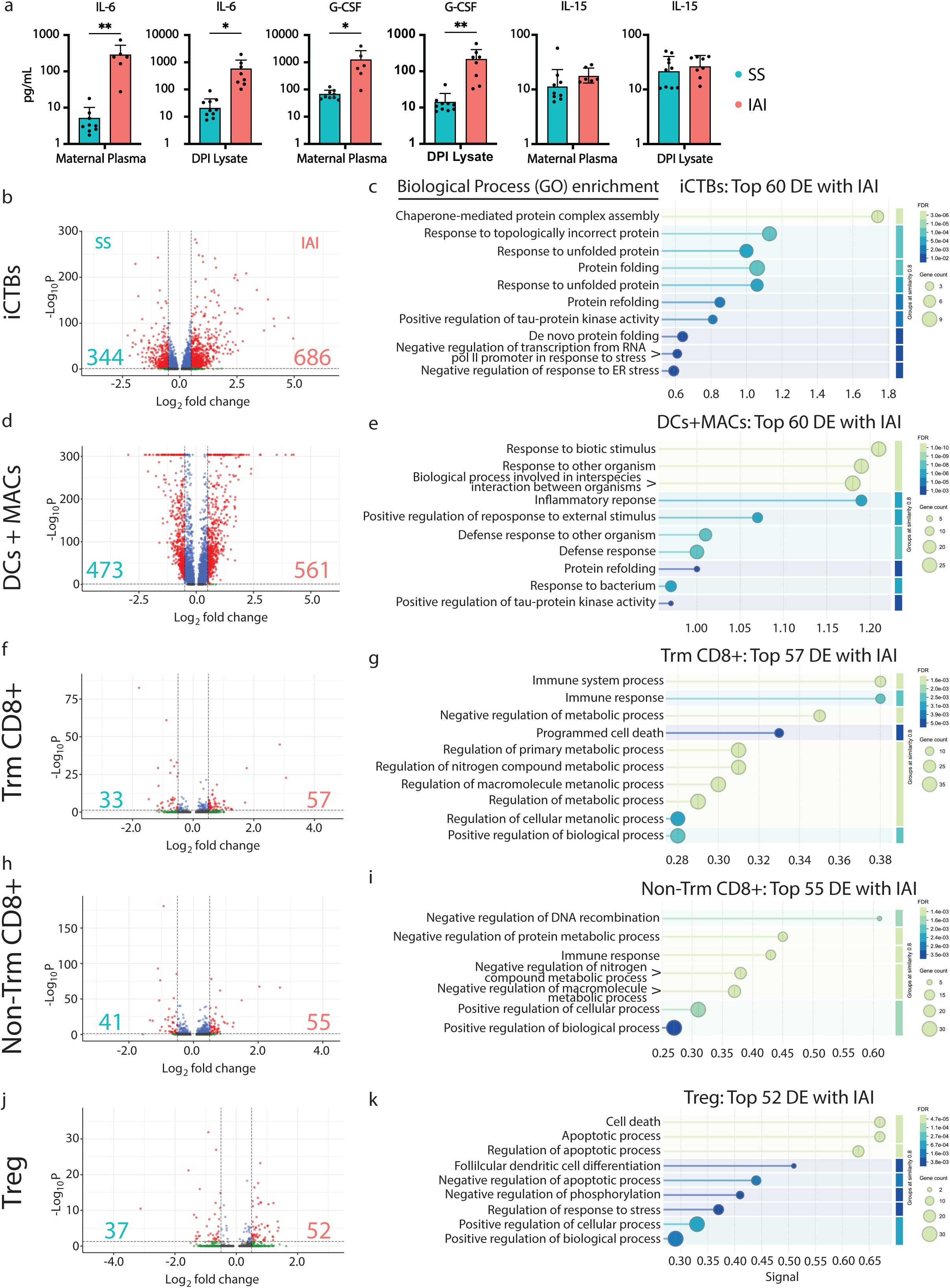
Intra-amniotic inflammation elicits a local and systemic inflammatory environment which does not activate T cells. **a,** Cyokine concentrations determined by multiplex analysis of maternal blood plasma or DPI lysate. Each point represents a study participant during SS or IAI; IL-6, G-CSF, and IL-15 within maternal blood plasma during SS (n=9) or IAI (n=6); IL-6, IL-15, G-CSF within DPI lysate during SS (n=10) or IAI (n=8). **b,d,f,h,j** Volcano plots depicting the differential gene expression between SS and IAI of iCTBs, dendritic cells and macrophages (APCs), Trm CD8^+^ T cells, Non-Trm CD8^+^ T cells, and Tregs. DE gene cutoff values were: adj P value=< 0.05, Log FC <-0.5 and >0.5. The genes above these cutoff ranges are depicted as red dots; the total number of red dots for each condition are printed in the bottom left or right of the plot, respectively. The top 10 DE genes for each condition are printed to the left or right of the plot, respectively. **c,e,g,I,k** Biological Process Gene Ontology enrichment analysis by StingDB. Error bars, SD. ** p<0.005; * p<0.05 by upaired T tests.

Akin to our SS samples, we used scRNAseq to examine cells from IAI placental tissue (n=6 donors; 26,807 cells) and the DPI (n=6 donors; 86,581 cells) (Supp Fig 2; Supp Fig 3). In our IAI samples we again observed a heterogeneous population of cytotrophoblasts divided into iCTBs and villous CTBs (Supp Fig 2 f). We also retrieved all key immune population from IAI tissues (Supp Fig 3). This allowed us to assess transcriptional changes between the two conditions within specific subpopulations.

We observed large number of transcriptional changes (over 1000 DE genes) between SS and IAI within the iCTB and APC populations (**Fig 3 b,d**), suggesting that both the tissue and APC populations robustly responded to inflammatory signals present during IAI. In contrast, we observed minimal transcriptional shifts within the Trm and non-Trm CD8^+^ T cells and Tregs, with less than 100 transcriptional changes between conditions (**Fig 3 f,h,j**).

To broadly classify the iCTB changes in response to IAI, we used StringDB to classify the top 60 DE genes increased with IAI (**Fig 3c**). Most of these Biological Processes were related to protein folding, suggesting a stress response by iCTBs within the IAI environment. Within the APC population, we found Biological Processes related to pathogen response and protein folding (**Fig 3e**). Despite their limited number of altered transcripts, we performed a StringDB analysis in Trm and non-Trm CD8^+^ T cells and Tregs which highlighted potential metabolic changes in the CD8 T cell subsets, and cell death-related changes in Tregs (**Fig 3g,i,k**).

We considered that the lack of T cell responses could be due to a lack of activating signals, the presence of inhibitory signals and/or a state of dysfunction (such as exhaustion or tolerance) of the CD8 T cells in the DPI.

### CD8 T cell subsets in the DPI are functional

To determine the phenotypic and functional properties of T cells isolated from the DPI, we used the flow cytometry dataset from a recent study(17) to assess the expression frequencies of PD1 (residency/inhibition/exhaustion), CD69 (activation), and GZMB (cytotoxic effector function) (**Fig 4a-d**).

**Figure 4:**
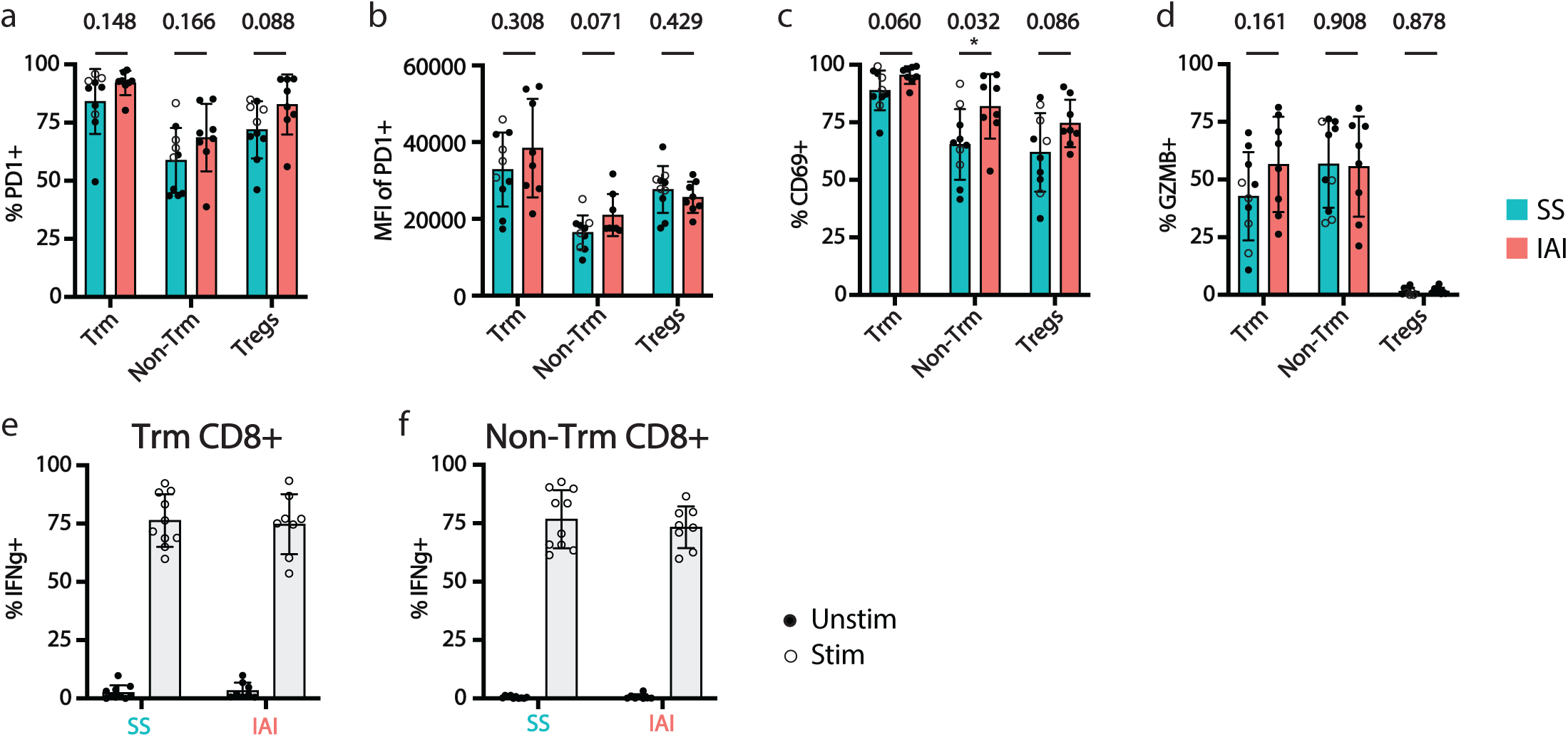
Although markers of T cell residency and activation are maintained during steady state and intra-amniotic inflammation, T cells are capable of effector function. **a-d,** We used a previously published flow cytometry dataset to assess frequencies and/or mean fluorescence intensity (MFI) of Programmed Cell Death 1 (PD1)^+^, CD69^+^, and Granzyme B (GZMB)^+^ within the Trm CD8^+^, Non-Trm CD8^+^, and Treg populations of the DPI. Each point represents a study participant during SS (n=10) or IAI (n=8). Labored samples in the SS condition are open circles. **e,f,** Frequencies of IFNg within the Trm CD8^+^ and Non-Trm CD8^+^ populations of the DPI during SS (n=10) or IAI (n=8) either ex vivo (Unstim) or following 6hr stimulation with PMA/Ionomycin (Stim). Error bars, SD. P values; * significant p<0.05 by unpaired T tests.

In T cells isolated from IAI DPI, PD1 protein expression did not change significantly across CD8 T cell and Treg populations from SS and IAI tissues (**Fig 4a**). Similarly, PD-1 expression levels per cell (MFI) remained unchanged as well (**Fig 4b**). Consistent with the modest signs of activation observed at the transcriptional level, we also find modest levels of increased CD69 protein expression, though only significantly within the non-Trm CD8^+^ T cells (**Fig 4c**). Lastly, we looked at effector function within these T cell subsets and find, in agreement with the scRNAseq data, the GZMB effector program for these T cells are maintained during health and IAI (**Fig 4d**).

To determine if the immune cells within the DPI are responsive to activating signals, we stimulated the immune cells with phorbol myristate acetate and ionomycin (PMA/Iono). We found that T cells isolated from SS and IAI tissues responded equally well to ex vivo stimulation, as observed by IFNγ production within the Trm (**Fig 4e**) and non-Trm CD8^+^ populations (**Fig 4f**). These results suggest that T cells from the DPI are not dysfunctional (as could be suggested by their expression of PD1) or tolerized (due to their residency within the semi-allogeneic placenta). Thus, the modest T cell activation in context of IAI did not indicate an intrinsic inability of T cells to respond to activating signals.

We next interrogated T cell – iCTB interactions during IAI. We used NicheNet to predict signals sent from from T cells to iCTBs in IAI samples (**Supp Fig 6a,b**). Here we found pathways integral to the survival and differentiation of iCTBs conserved among all T cells such as: BMP7/ACVR1, BMP7/ACVR2A(36), TRAIL/DR5(37, 38), CSF1/CSF1R(26), CCL5/CCR1(26), LTB/LTBR(38), NRG1/ERBB3(39), TGFβ/ TGFβR1(25), TGFβ/ TGFβR2(25), CCL5/CCR3, SEMA4D/PLXNB1(40, 41); possible immune modulation: IL1B/IL1R1(42), IL1B/IL1R2(42), IFNG/IFNGR1, IFNG/IFNGR2(43, 44), PTPRC/CD22^25^. Looking at predictions shared by specific T cell subsets, we find the following pathways integral to trophoblast survival and differentiation shared by nonTrm and Trm to iCTBs: XCL1/XCR1(26), LIGHT/TLBR(38), VEGFB/FLT1(45–47); shared by nonTrm and Tregs to iCTBs: TNF/TNFRSF1A(48), CCL4/CCR1(49); and shared by Trm and Tregs to iCTBs: LTA/LTBR(38). Together, these data indicate that T cells maintained their potential to support iCTB homeostasis during a state of acute tissue inflammation (IAI). This outcome was somewhat expected given the limited transcriptional changes observed for T cell subsets isolated from IAI tissues.

We next used MultiNicheNet to identify the top 25 T cell – iCTB interactions that differ between SS and IAI, then examined ligand activity for these interactions. We observed some changes in signals related to iCTB survival and differentiation (bolded in **Fig 5a**), including CCL5 and TRAIL (**Fig 5a**). We next validated the presence of predicted key iCTB survival and differentiation factors within the DPI environment. Within DPI lysates, we measured the protein level of CSF1, CCL5, XCL1, and TRAIL and found their overall levels are consistently maintained within the DPI environment between SS and IAI conditions (**Fig 5b**). Overall, these data do not exclude the possibility that some T cell-derived signals changed during IAI, but also suggest that these changes did not profoundly affect the availability of these signals in the DPI. Given the substantial transcriptional changes of iCTBs during IAI, we next asked if iCTBs maintained their ability to recruit and maintain T cells during IAI.

**Figure 5:**
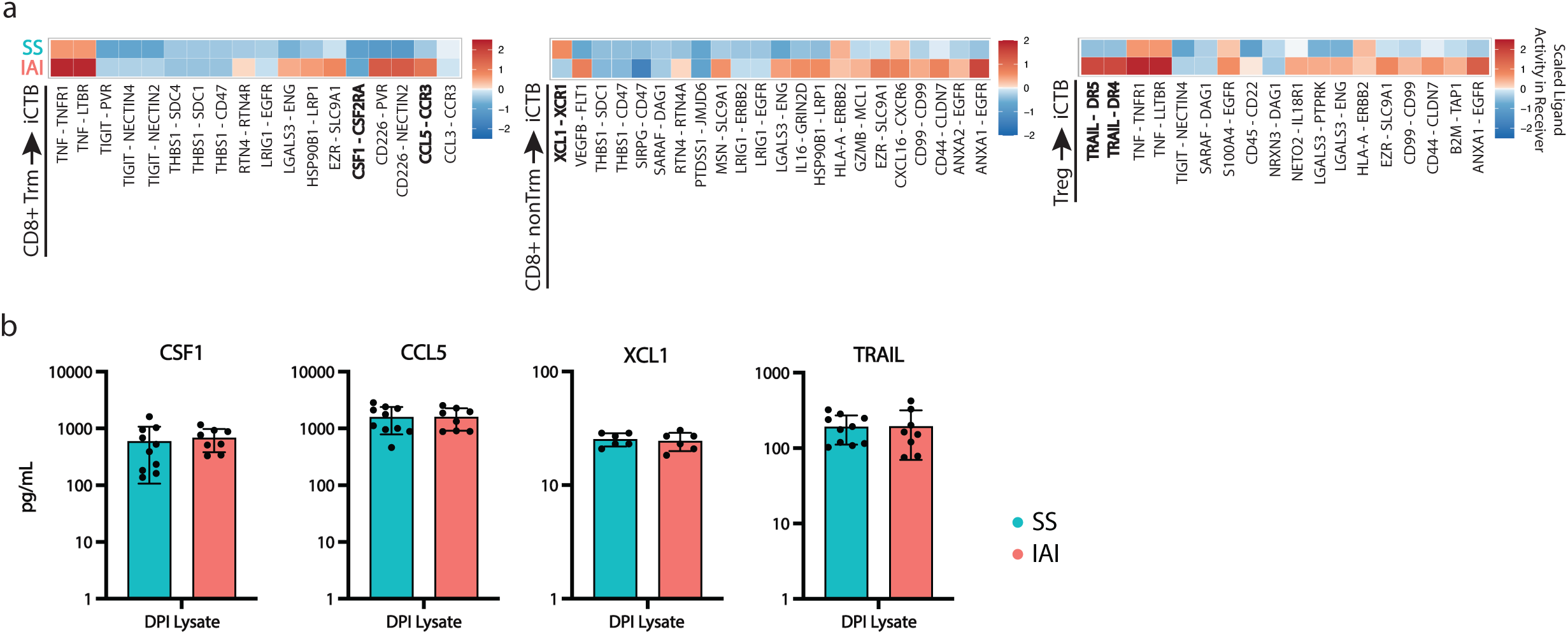
Support for trophoblast survival and differentiation is maintained during IAI. **a,** Plots visualizing the Ligand Activity in Receiver cells of the top 25 MultiNicheNet-predicted interactions between T cells and iCTBs during IAI **b,** Cyokine concentrations determined by multiplex analysis of DPI lysate during SS or IAI. Each point represents a study participant; CSF1, CCL5, TRAIL during SS (n=10) or during IAI (n=8); XCL1 during SS (n=6) or during IAI (n=6). Error bars, SD.

### iCTB maintain their support for CD8 T cell recruitment during IAI

We initially performed a NicheNet analysis for iCTB signals sent to T cells for the IAI condition. We found predicted signals from iCTBs to T cells related to the adhesion and chemotaxis of T cells: CCL4/CCR5(50), CXCL16/CXCR6(51, 52), and possible immune modulation PVR/CD226, PVR/TIGIT, NECTIN3/TIGIT(53), and signals related to survival and differentiation: TGFβ1/ TGFβR2, PD-L1/ PDCD1 (30), PD-L2/PDCD1, TNFSF10/TNFRSF10B, TNF/TNFRSF1B, IGF2/IGF2R, EBI3/IL27RA (54). This analysis suggested that iCTBs maintained their potential to support T cell recruitment and homeostasis,

To assess how these interactions changed between SS and IAI, we again used MultiNicheNet. Looking at the ligand activity within the top 25 MultiNicheNet-predicted interactions between iCTBs and T cells, we observed a predicted increase in signaling supporting Trm residency, survival, and differentiation (e.g. PD-L1-PD1) (**Fig 6a**).

**Figure 6:**
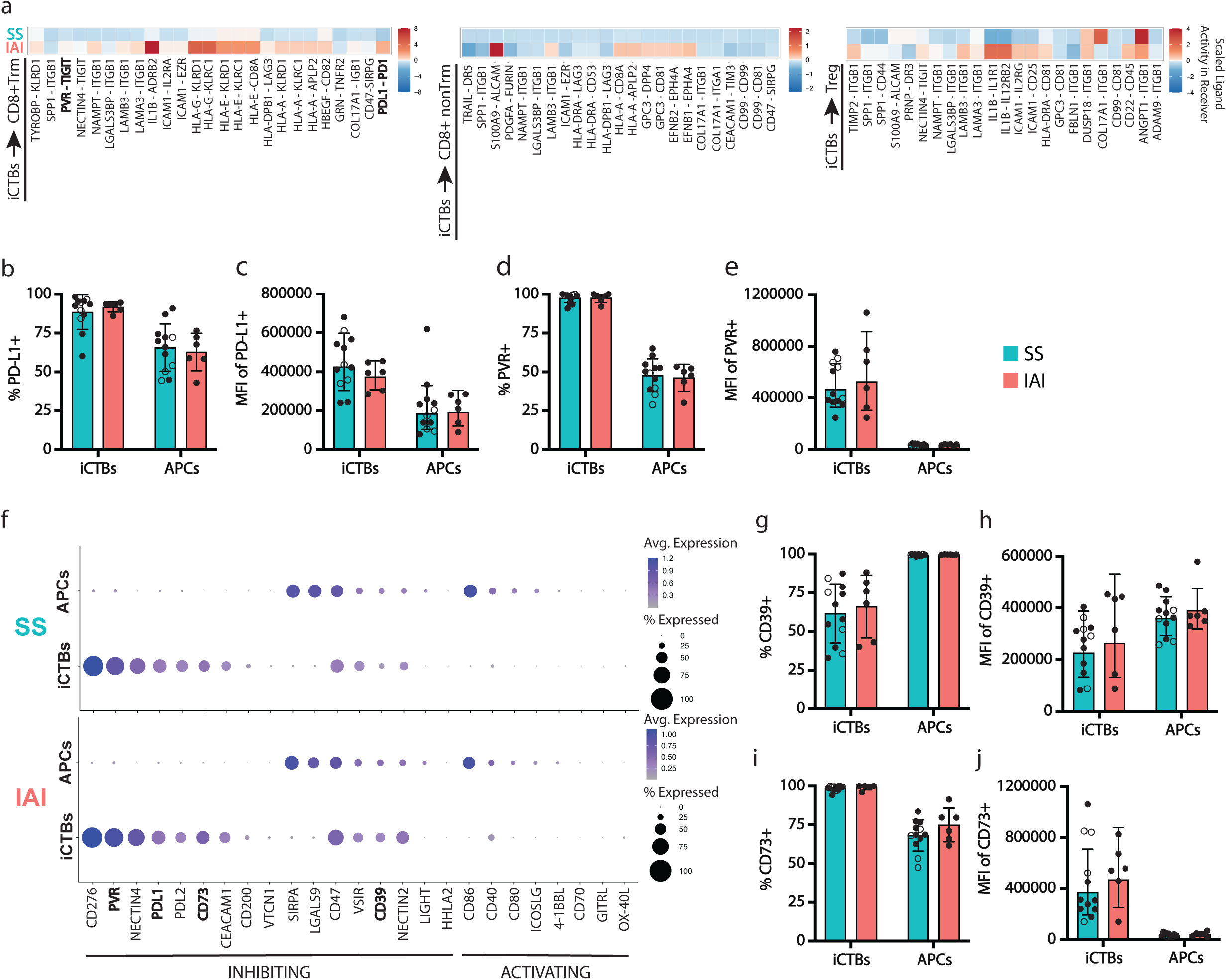
In addition to maintaining support for Trm survival and homeostasis, iCTBs may limit T cell activation. **a,** Plots visualizing the Ligand Activity in Receiver cells of the top 25 MultiNicheNet-predicted interactions between iCTBs and T cells during IAI. **b,c,d,e,** Frequencies and/or MFI of PD-L1 and PVR within the iCTB and APC populations during SS (n=12) and IAI (n=6). **f,** Bubble plot depicting average protein expression level and frequency of inhibitory or activating T cell signals by iCTBs or APCs during SS or IAI. Genes of interest with corresponding flow cytometry analysis are bolded. **g,h,i,j,** Frequencies and/or MFI of CD39^+^ and CD73^+^. Error bars are standard deviation from the mean. Labored samples in the SS condition are represented by open circles.

We next examined some predicted interactions by flow cytometry to validate predicted interactions between iCTBs and T cells during SS and IAI on the protein level (**Fig 6**; Supp Fig 7; Supp Fig 7). We found the frequency of PD-L1^+^ iCTBs and the MFI of PD-L1^+^ signal among iCTBs (**Fig 6b,c**) was maintained between health and IAI. We also examined the frequency and MFI of PD-L1 expression among APCs and found lower PD-L1 expression in comparison to iCTBs (**Fig 6b,c**). These data suggest that the iCTBs could play a key role in facilitating PD-1 dependent T cell tissue residency (29, 30), and controlling T cell reactivation within the DPI. Along with supporting T cell residency within the DPI, the expression of PVR by iCTBs was predicted to engage both T cell activating and inhibitory pathways via CD226 (DNAM1) and TIGIT, respectively (**Fig 2c**; **Fig 5a**; **Fig 6 a**). Of note, iCTBs almost uniformly expressed PVR (**Fig 6d**), and the expression levels (MFI) were several log fold higher compared to APCs (**Fig 6e**). Upon finding that iCTBs and APCs exhibit significantly different PD-L1 and PVR expression profiles, we looked across several other signals involved in T cell inhibition and activation to further dissect the immunoregulatory division of labor between iCTBs and APCs within the DPI. We found iCTBs and APCs exhibit an inverse distribution of functional roles at the transcript level across most markers, suggesting they may occupy functionally complementary niches within the DPI (**Fig 6f**). Consistent throughout the current study, these functional niches were maintained during both health and IAI. We also observed this pattern for CD39 and CD73, two cell surface proteins responsible for hydrolyzing pro-inflammatory ATP to adenosine. Unlike what we observed with PD-L1 and PVR where iCTBs were the main expressors, APCs expressed CD39 at a higher frequency and MFI than the iCTBs (**Fig 6g,h**). CD73 expression, however, was nearly uniform on iCTBs (**Fig 6i**) and several log fold higher on iCTBs that APCs (**Fig 6j**). These expression differences were significant between populations and were maintained between SS and IAI.

Overall, our data suggest that iCTBs and T cell communication is largely maintained during inflammation, which could ensure mutual support for cell survival and differentiation.

## DISCUSSION

Several studies have demonstrated the ability of regulatory T cells (Tregs) to directly support the survival and differentiation of tissue cells during steady state (SS), such as regulating hair follicle stem cell proliferation (15, 55–58). Taken together, these studies illustrate the function of Tregs beyond immune regulation and highlight their role in maintaining proper tissue function (59, 60). Tissue-resident memory T cells (Trm) are thought to surveil tissue sites (11) to provide localized pathogen protection (8, 61, 62). If and how Trm contribute to tissue cell survival and differentiation is less explored. Since trophoblasts do not express classical MHC class I molecules that would allow CD8 T cells to recognize antigen in an MHC-restricted manner, we initially asked if CD8 T cells in the decidual-placental interface (DPI) could provide any tissue-supporting functions.

Of note, we examined CD8 T cells within the third trimester DPI during SS and intra-aniotic inflammation (IAI). T cells increase in abundance in the placenta during the third trimester, while natural killer (NK) cells are the most prevalent immune cell type in the first and second trimester (63). Previous studies looking earlier in gestation have found NK cells to be integral to tissue remodeling, trophoblast differentiation, and invasion (26, 64, 65). Whether T cells can similarly support trophoblasts has not been previously investigated to the best of our knowledge, and it was also unclear how T cell – trophoblast communication may change in the context of acute inflammation.

While we initially intended to focus on communication of CD8^+^ Trm with trophoblasts, we found that many signals sent by CD8 Trm to iCTBs were also shared by non-Trm, and Tregs. Overall, we found that most predicted interactions between Trm and iCTBs were indicative of a mutual support for survival and differentiation. Furthermore, this is maintained during both SS and IAI across several predicted signaling exchanges. The differentiation of both Trm and iCTBs depends on TGFβ (25, 66–68), a stable prediction between SS and IAI. In addition to TGFβ, Trm formation and maintenance appears to also depend on PD-L1-PD1 signaling (30). Our data indicate that iCTBs provide stable PD-L1 signaling to Trm. Importantly, when we examined PD-L1 expression on the protein level, iCTBs were the main expressors of PDL1 within the DPI, expressing a significantly higher frequency and mean fluorescence intensity (MFI) compared to the APCs. during both SS and IAI, we hypothesize iCTBs recruit (CXCL16-CXCR6 (51, 52)) and support (TGFβ,PD-L1) Trm for the number of survival and differentiation signals they provide. Of note, Trm were predicted to provide several factors important for iCTB survival and differentiation, during both SS and IAI. Factors such as CCL5, XCL1, CSF1, produced by uterine NK cells during early pregnancy, regulate trophoblast differentiation (26). Here we show Trm were predicted to provide these signals to iCTBs at full-term, during both SS and IAI. It is important to consider that IAI is associated with eliciting labor, which may further affect phenotype and function of immune cells in the DPI. Although dissecting these signals is not the goal of this study, we included samples from labored healthy individuals for flow cytometry protein validation in an effort to control for this variable.

Beyond providing survival and differentiation support, we find that while Trm did not increase their activation status or GZMB expression with IAI, they were functional ex vivo as indicated by IFNγ^+^ production upon activation. This suggests that Trm within the DPI can provide protection from pathogens, although acute inflammation in the form of IAI did not elicit acquisition of effector function, potentially to avoid bystander-activated CD8^+^ T cell-mediated tissue injury (69). Overall, our data show that IAI and the associated increase in local and systemic IL-6 elicited substantial transcriptional changes in iCTBs and APCs, but only elicited very limited changes in Trm and other T cell subsets. Whether this is due to a lack of needed activating signals or the presence of inhibitory signals within the DPI is beyond the scope of this investigation.

Lastly, our data suggest a “division of labor” relationship between iCTBs and APCs within the DPI. Looking across many markers (at the transcript level) known to play a role in the inhibition or activation of T cells, we find that markers expressed by iCTBs were expressed to a lesser extent by APCs, and vice versa. Furthermore, this is maintained during both SS and IAI. Importantly, expression patterns on the transcript level may not translate to the protein level(70). However, for the select markers that we examined at the protein level, we were able to validate significantly different levels of expression between iCTBs and APCs. Taken together, our data suggest a robust communication network between tissue (iCTBs) and T cells that is largely maintained during acute inflammation, and could serve to ensure that acute inflammatory events are not sufficient to elicit harmful events at the decidual-placental interface.

## METHODS

### Sex as a Biological Variable

This study investigated tissue-immune interactions within the human placenta and therefore included blood and placenta tissue samples from consenting pregnant females.

### Diagnostic criteria

Study participants were diagnosed with intra-amniotic inflammation (IAI) if they had a temperature above 38°C with maternal and/or fetal tachycardia. H&E-stained placenta tissues from subjects at SS or with IAI were evaluated for inflammation by a blinded gynecologic and breast pathologist from the University of Washington Department of Laboratory Medicine and Pathology.

### Tissue mounting and Immunofluorescence

Tissue samples spanning the full-depth of the placenta, including from the decidua basalis to the fetal membranes, were collected using a fresh regular duty single edge razor blade to cut through the placenta, taking care not to twist or crush the tissue. As previously described(71), excised tissues were rinsed in PBS and fixed overnight at 4°C in Cytofix buffer (BD Biosciences 554655) diluted 1:4 in PBS. Fixed tissues were next dehydrated overnight at 4°C in sterile-filtered 20% w/v sucrose (Sigma Aldrich). Tissues were then mounted in Tissue-Tek OCT medium (Sakura Finetek) in Peel-A-Way disposable plastic tissue embedding molds (Polysciences, Inc.) and flash frozen in the vapor phase of LN_2_ and stored at −80°C.10-12μm sections of tissue were cut using a CM1950 cryostat (Leica), dried overnight at room temperature (RT), and stored at - 80°C. When ready to stain, slides were incubated in −20°C acetone for 5 min (in glass coplin jar), after which excess acetone was wiped off, and tissue was outlined with a hydrophobic pap pen (Vector Laboratories H-4000). Slides were rehydrated in 1x TBS-T for 5 minutes at RT (in plastic coplin jar), then tissues were blocked with 50μl normal human serum (Fisher Bio 31876) (1:20 Normal Human Serum:1x TBS-T) for 30 min. Primary unconjugated antibodies, mouse anti-human Cytokeratin 7 (CK7) (Agilent M701829-2; clone OV-TL 12/30; 1:100) and rabbit anti-human Vimentin (Vim) (Abcam ab16700; clone SP20; 1:1000), were incubated for 60 minutes at RT or overnight at 4°C in 50μL blocking buffer, rinsed for 5 minutes in TBS-T (in plastic coplin jar), then incubated with secondary conjugated antibodies in 50μL blocking buffer for 60 minutes at RT. AF594-conjugated anti-mouse IgG (Life Technologies A11001; 1:200) was used for CK7 and AF488-conjugated anti-rabbit IgG (Life Technologies A31573; 1:200) was used for Vim. Prior to staining tissues with the conjugated CD8 antibody, tissues were blocked with 50μL normal mouse serum (Thermo 10410) (1:20 Normal Mouse Serum:1x TBS-T) for 60 minutes at RT, after which, the slides were incubated with AF647-conjugated CD8 antibody (BD 557708; clone RPA-T8), and rinsed for 5 minutes in TBS-T. Tissues were incubated in 50μL DAPI (ThermoFisher F10347) (1:1000 in PBS) for 5 minutes then rinsed in PBS for 5 minutes. To reduce tissue autofluorescence the following steps were performed in coplin jars: slides were dunked in ammonium acetate, incubated in copper sulfate for 5 minutes, dunked in ammonium acetate, dunked in PBS, then dunked in dH_2_O. Cover slips were mounted using 20-50μL of Prolong Gold antifade reagent (Invitrogen P36930) and allowed to cure for 24 hours.

### Transmission Electron Microscopy

Placental tissue was fixed in half-strength Karnovsky’s fixative (2.5% glutaraldehyde, 2% paraformaldehyde in 0.1M sodium cacodylate buffer, pH 7.3) overnight at 4 degrees C. Tissue pieces were rinsed with 0.1M cacodylate buffer, treated with 1% osmium tetroxide for 2 hours (4 degrees C), rinsed with cacodylate buffer and dehydrated through a graded series of ethanols and propylene oxide and embedded in Eponate12 resin (Ted Pella, Inc, Redding, CA). 70nm ultrathin sections were cut using a Leica EM UC7 ultramicrotome, contrasted with uranyl acetate and lead citrate and imaged on a ThermoFisher Talos L120c transmission electron microscope at 120kV. Digital images were acquired with a Ceta 16M CMOS 4kx4k digital camera system. Light level 1 micron sections were stained with filtered Richardson’s methylene blue (10% methylene blue, 10% azure II, 10% sodium borate) for examination. This research was supported by NIH P30 CA015704 of the Fred Hutch/University of Washington/Seattle Children’s Cancer Consortium, which includes the Electron Microscopy Shared Resource.

### Tissue Digestion and single cell isolation

All samples were maintained on ice following acquisition and best efforts were made to initiate tissue digestion within two to four hours of delivery. Sterile instrumentation was used for all tissue processing.

#### Isolation of cytotrophoblasts from placentas

Cytotrophoblasts (CTBs) were isolated from full-term placentas as previously described (72). In brief, CTBs from the floating villi and the decidual-placental interface (DPI) were isolated in the following manner. Snippets of tissue sampled from throughout the placenta were placed in ice-cold PBS and minced to 1mm^3^ pieces. Minced tissue was washing in PBS to remove blood. Minced tissue was digested using first a collagenase digest to remove the syncytiotrophoblast (18.2 units/100mL collagenase (Sigma C2674-1G), 0.012g/100mL DNAse (Roche 104159), 0.1g/100mL bovine serum albumin (Sigma A7906-100G), and 0.069g/100mL hyaluronidase (Sigma H3506-5G) in PBS), shaking at 37°C for 40 minutes. At the end of 40 minutes, the undigested tissue was allowed to settle and the supernatant containing the digested syncytiotrophoblast was discarded. Two subsequent DNAse digestions were performed to isolate cytotrophoblasts (20mL/100mL Trypsin with EDTA (Gibco 25200-056), 80mL/100mL PBS, 0.012g/100mL DNAse (Roche 104159)), the first shaking at 37°C for 10 minutes and the second for 5 minutes. A final rinse with cytowash media (2.5% FBS, 1% penicillin streptomycin (Gibco 15140-122), 1% Gentamycin (Sigma-Aldrich G1397-10ML), 1% GlutaMax (Gibco 35050-061)in DMEM (Gibco 11965)) was used to collect remaining digested cells. After each digest, the undigested tissue was allowed to settle, and the digested cells were poured over fresh sterile gauze sponges into a 500mL glass bottle containing 50mL FBS for collection. Single cell digests were pooled. Digest volumes were calculated as follows: 6 x # of grams of tissue for the collagenase and first DNAse digests, and 5 x # of grams of tissue for the second DNAse digest. The pooled single cell digest was spun down at 314*g* for 10 min and pellets were resuspended, pooled in 40mL of cytowash, and filtered using a 100m cell strainer (Corning 352360). The filtered cells were spun down at 314*g* for 10 min and pellets were resuspended in 4mL of cytowash per 7g digested tissue. To enrich for CTBs, 4mL of tissue digest was layered onto a six layer percoll (Sigma-Aldrich P1644-500ML) gradient (70%-20%) and spun at 1589*g* for 25 min. Layers containing 30 and 40% percoll were isolated and enriched CTBs were washed in culture medium via centrifugation (314*g* for 10 min). Single cell digests were resuspended in Recovery Cell Culture Freezing Medium (Gibco 12648-010) at a concentration of 1-5×10^6^ cells/mL.

#### Isolation of leukocytes from placentas

Leukocytes were isolated from the decidual-placental interface (DPI), which includes the decidua basalis, of full-term placentas as previously described(71). In brief, 0.5cm-depth biopsies were collected from the DPI. Tissues were minced in digestion media: RP7.5 + 700U/mL Collagenase Type II (Sigma Aldrich), 200U/mL DNase (Sigma Aldrich), and incubated in a shaking 37°C incubator for 60 min at 225 RPM (). Following digestion, single-cell suspensions were strained through a 70µm filter, cells were centrifuged at 400*g* for 5 min, and washed with RP7.5 media. Red blood cells were removed using ACK lysis, and cell pellets were resuspended in RP7.5 before downstream use. Guava easyCyte was used to quantitate cells. Single cell digests were resuspended in cryopreservation medium (10% sterile DMSO (Sigma Aldrich), 90% FBS).

#### Isolation of plasma and leukocytes from blood

Maternal peripheral blood (mBlood) was collected prior to delivery and fetal cord blood (cBlood) immediately after delivery by venipuncture as previously described(71). In brief, blood samples were collected in acid citrate dextrose solution A (ACD)-vacutainer tubes (BD Biosciences). Blood was centrifuged at 400*g* for 10min and plasma was collected and stored at −80°C.

Leukocytes were isolated via ACK lysis (ThermoFisher) and then processed using SepMate tubes (StemCell Technologies, 85450) and Lymphoprep (Stem Cell Technologies, 07851) according to manufacturer protocols. In brief, remaining cells were resuspended in 30 ml PBS and pipetted on top of 13.5 ml Lymphoprep in a SepMate tube. After centrifugation for 16 min at 1,200*g*, the mononuclear cell fraction in the supernatant was poured into a fresh 50-ml tube, washed with PBS. Cells were resuspended in cryopreservation medium (10% sterile DMSO (Sigma Aldrich), 90% FBS) and stored in liquid nitrogen until used for downstream procedures.

### scRNAseq

scRNAseq libraries generated from DPI, mBlood, and cBlood immune cell isolations were generated following the Rhapsody BD Rhapsody^TM^ System TCR/BCR Full Length, mRNA Whole Transcriptome Analysis (WTA), BD AbSeq, and Sample Tag Library Preparation Protocol. Libraries generated in 2022 followed version 23-24020(01) and libraries generated in 2024 followed version 23-24020(02). scRNAseq libraries generated from CTB cell isolations were generated following the Rhapsody BD Rhapsody^TM^ System mRNA Whole Transcriptome Analysis (WTA) and Sample Tag Library Preparation Protocol version 23-24119(02).

#### Whole Transcriptome single-cell library preparation and sequencing

cDNA libraries were generated as previously described (73). In brief, after sorting and removing dead cells (see Flow Cytometry and Cell Sorting section below), single cells were stained with Sample-Tag antibodies, stained with AbSeq antibodies (DPI/mBlood/cBlood samples only) as previously described (73), washed three times, pooled and counted and subsequently loaded onto a nano-well cartridge (BD Rhapsody), lysed inside the wells followed by mRNA capture on cell capture beads according to manufacturer instructions. Cell Capture Beads were retrieved and washed prior to performing reverse transcription and treatment with Exonuclease I. cDNA underwent amplification via PCR (10–11 cycles). PCR products were purified, and mRNA PCR products were separated from Sample-Tag (and AbSeq, where applicable) PCR products with double-sided size selection using SPRIselect magnetic beads (Beckman Coulter). mRNA and Sample Tag products were further amplified using PCR (10 cycles). PCR products were then purified using SPRIselect magnetic beads. Quality of PCR products was determined by using an Agilent 2200 TapeStation with High Sensitivity D5000 ScreenTape (Agilent) in the Fred Hutch Genomics Shared Resource laboratory. The quantity of PCR products was determined by Qubit with Qubit dsDNA HS Assay (Q32851). mRNA product was diluted to 2.5 ng μl^−1^, and the Sample Tag and AbSeq PCR products were diluted to 1 ng μl^−1^ to prepare final libraries. Final libraries were indexed using PCR (6 cycles). Index PCR products were purified using SPRIselect magnetic beads. Quality of all final libraries was assessed by using Agilent 2200 TapeStation with High Sensitivity D5000 ScreenTape and quantified using a Qubit Fluorometer using the Qubit dsDNA HS Kit (ThermoFisher). Final libraries were diluted to 3 nM and multiplexed for paired-end (100 bp) sequencing on a NovaSeq 6000 (Illumina) using S1 and S2 flow cells. For the gene expression library, we targeted 5,000–20,000 reads per cell, for the AbSeq library 10,000–15,000 reads per cell, and for the Sample-Tag libraries 500–2,000 reads per cell.

### scRNAseq Analysis

Using the Seven Bridges Genomics website, we used the BD Rhapsody Sequencing Analysis Pipeline (Revision 15) to align FASTQ files with a genome reference file (GRCh38 Human Rhapsody Reference specified by BD for use with their Sequencing Analysis Pipeline) as well as an AbSeq reference file where applicable; the output of which were aligned Seurat objects. The R package, Seurat 5.1.0 (74), was used for downstream analysis.

#### Trophoblast cell selection for downstream analysis

All scRNAseq libraries generated from cytotrophoblast isolations (including both SS and those with IAI) were combined into a single Seurat object and batch corrected using Harmony. A combination of transcripts was used to identify the trophoblast-containing clusters: KRT7^+^, HLA-G^+/-^, EGFR^+/-^, MKI67^+/-^, VIM^-^, CD45^-^, CD14^-^, CD19^-^, CD3E^-^ (Supp Fig 1). The trophblasts were further purified by keeping cells that expressed CD45 (transcript <1) and VIM (transcript <1) (Supp Fig 1 c). Cells labeled as “Mulitplet” and “Undetermined” were removed. The purified trophoblast population was rescaled, reharmonized, and re-clustered and the invasive cytotrophoblasts (HLA-G^+^) were used for downstream analysis (Supp Fig 1). Data was subsetted by disease condition (SS or IAI) when applicable.

#### Immune cell selection for downstream analysis

All scRNAseq libraries generated from cells isolated from cBlood, mBlood, and the DPI (including both SS and those with IAI) were combined into a single Seurat object and batch corrected using Harmony. Cells labeled as “Mulitplet” and “Undetermined” were removed. A combination of transcript expression and antibody-sequencing (AbSeq) was used to determine immune cell lineages (Supp Fig 2 c). The following AbSeq lineage markers were used to identify the <u>memory CD8+ T cell-containing cluster:</u> CD3^+^, CD45RO^+^, CD45RA^lo^, CCR7^lo^, CD4^lo^, CD8^+^, CD14^-^ (Supp Fig 2 c). After subsetting, rescaling, reintegrating (Harmony for batch correction), and re-clustering, this object was separated by cBlood, mBlood, and DPI to determine if cells from the DPI samples were also present in the cBlood and mBlood samples, indicating “blood contamination” (Supp Fig 3 a). Clusters which were unique to the DPI sample were subsetted and considered “DPI -specific” memory CD8 T cells. The memory CD8 T cells were further purified by keeping cells that expressed TCRgd (AbSeq <2), SLC4A10 (transcript <1; MAIT cell marker), and CD4 (AbSeq <1) (Supp Fig 3 c). Finally, DPI -specific memory CD8 T cells were separated into Trm and non-Trm populations based on CD103 expression (AbSeq <>2) (Supp Fig 3 c) and used for downstream analysis. FOXP3 transcript expression was used to identify the <u>Treg population</u> (Supp Fig 2 c). Like above, after subsetting, rescaling, reintegrating (Harmony for batch correction), and re-clustering, this object was separated by cBlood, mBlood, and DPI to determine if cells from the DPI samples were also present in the cBlood and mBlood samples, indicating “blood contamination” (Supp Fig 3 e). Cells which were unique to the DPI sample were subsetted and considered “DPI -specific” Treg cells and used for downstream analysis. The AbSeq lineage marker CD14 was used to identify the <u>APC cluster</u> (Supp Fig 2 c). Like above, after subsetting, rescaling, reharmonizing, and re-clustering, this object was separated by cBlood, mBlood, and DPI to determine if cells from the DPI samples were also present in the cBlood and mBlood samples, indicating “blood contamination” (Supp Fig 4 a). Cells which were unique to the DPI sample were subsetted and considered “DPI-specific” APC cells and used for downstream analysis. Data was subsetted by disease condition (SS or IAI) when applicable.

The following parameters were applied to otherwise standard NicheNet(75) (v2.1.5) and MultiNicheNet (v2.0.0) analyses. A gene was considered expressed if it is expressed in at least 10% of cells in a population. The top 100 ligands were considered when calculating the best upstream ligands. NicheNet was applied to predicted ligand-receptor interactions between select tissue-immune populations within either the SS or IAI condition, using markers of those populations. MultiNicheNet was applied to predict ligand-receptor interactions between select tissue-immune populations based on the differential expression profiles between SS and IAI. The top 25 NicheNet-predicted interactions were visualized by Circos plot (Fig 2) and the top 25 MultiNicheNet-predicted interactions were visualized as “ligand activity in receiver” (Fig 5,6).

### Flow/FACS

#### Flow cytometry and cell sorting

For flow cytometric analysis good practices were followed (76). Following isolation or thawing, cells were wash 1 time in PBS and then incubated with Fc-blocking reagent (BioLegend Trustain FcX, 422302) and fixable UV Blue Live/Dead reagent (ThermoFisher, L34961) in PBS (Gibco, 14190250) for 15 min at RT. After this, cells were incubated for 20 min at RT with 50 μl total volume of antibody surface stain freshly prepared in Brilliant staining buffer (BD Biosciences, 563794), followed by two washes in fluorescence-activated cell sorting (FACS) buffer (PBS with 2% FBS). All antibodies were titrated and used at optimal dilution, and staining procedures were performed in 96-well u-bottom plates (for cell sorting in 5-ml polystyrene tubes). Panels were designed according to best practices as described(77). For sorting, cells were immediately used after staining. For analysis the stained cells were fixed with 4% PFA (Cytofix/Cytoperm, BD Biosciences, 554722) for 20 min at RT. If necessary, intracellular or intranuclear staining was performed following the appropriate manufacturer protocols (eBioscience FOXP3/Trascription Factor Staining Buffer Set, Thermo Fisher 00-5532-00). Cells were then washed, resuspended in FACS buffer and stored at 4°C in the dark until acquisition.

Single-stained controls were prepared with every experiment using antibody capture beads (BD Biosciences anti-mouse (552843) or anti-mouse Plus, and anti-rat (552844)) or cells where appropriate, and treated the same as the samples (including fixation procedures). For each staining of experimental samples, a PBMC was stained with the same panel as a longitudinal reference control.

Data aquisition for flow cytometry analysis was performed on a FACSDiscover S8 Cell Sorter (BD Biosciences) and FACSDiva acquisition software (BD Biosciences). Full details on the optical configuration of the instruments used are as described(78). After acquisition, data was exported in FCS 3.1 format and analysed using FlowJo (version 10.10.x, BD Biosciences). Samples were analysed using a combination of manual gating and computational analyses approaches(76). Gates were kept the same across all samples except where changes in the density distribution of populations clearly indicated the need for sample-specific adjustment. We required a minimum number of 25 cells for downstream analysis; all samples met this requirement (Supp Fig 7 c).

All lymphocyte cell sorting was performed either on a FACSAria III (BD Biosciences) or a FACSDiscover S8 Cell Sorter (BD Biosciences). Samples from four participants were enriched for specific immune cell subsets as previously described(79).

### Dead cell removal from single cell cytotrophoblast isolation

Dead cells were removed from cytotrophoblast (CTBs) isolations for immediate downstream single cell RNA sequencing processing either by sorting on a MACSQuant Tyto Cell Sorter (Miltenyi) into cytowash medium or with the Miltenyi Dead Cell Removal Kit (130-090-101) with Miltenyi LS Columns (130-042-401), following manufacturer’s instructions.

### Cytokine and chemokine quantification

Cytokine and chemokine quantification was performed on tissue lysates and blood plasma samples. To obtain lysates from placenta tissues, multiple 0.5cm-depth samples from the DPI were minced in lysing buffer (PBS/0.05% Tween 20 (ThermoFisher)) at 1g tissue/mL buffer and then centrifuged at 10,000 rpm for 5 min at 4°C. The supernatant was collected and immediately stored at −80°C. Tissue lysate supernatants and plasma were sent to Eve Technologies for protein quantification.

### Ex vivo Stimulation Assay

Cells were isolated from tissues or blood as described above. 1-2 million cells were placed into each well of a U-bottom 96-well plate with 200 μl R10. Cells were then left untreated (control) or stimulated with PMA (50 ng/ml) and ionomycin (500 ng/ml) for 6 h at 37 °C. Cells were then washed with 1× PBS prior to antibody staining and downstream flow cytometry analysis.

### Statistics

Unpaired, two-tailed, T tests were performed on flow cytometry and cytokine data to assess significance between SS and IAI conditions. Significance was indicated by an *.

### Study population and approval

Samples were obtained from pregnant individuals at the University of Washington Medical Center (Seattle, WA) under the Institutional Review Board (IRB) approved study: STUDY00001636. All participants gave signed consent prior to participation. Participants were over the age of 18, delivering full-term (37-41 weeks gestation) singleton pregnancies by either cesarean or vaginal delivery. Individuals with the following pregnancy complications were excluded: multiple gestation, preeclampsia, preterm labor, or any placental abnormalities (e.g., previa, abruption, etc.).

### Data Availability

Single cell RNA sequencing data is available on GEO, Accession # GSE327349. Flow cytometry data shown in Figs. 1 and 4 are from a previous study, and have been deposited on ImmPort (SDY3484). Flow cytometry data from Fig. 6 are available upon request.

## ACKNOWLEDGEMENTS

The authors thank the University of Washington Medical Center Labor and Delivery staff, study participants, and research coordinators for their participation in this study. We thank the laboratory of Susan J. Fisher at the University of California, San Francisco, for hosting C.S.D. to learn methods in placental biology. We thank Daniel J. Reiter, M.D., a gynecologic and breast pathologist at the University of Washington Medical Center Department of Laboratory Medicine and Pathology for evaluating placental inflammation. We thank Fred Hutch Cancer Center Shared Resources including the Electron Microscopy, Imaging, Genomics, Experimental Histopathology, and Flow Cytometry cores for technical assistance.

## AUTHOR CONTRIBUTIONS

Research study design (C.S.D., S.A.M., M.P.), conducting experiments (C.S.D., M.F., N.B.P., G.D., A.J.K.), acquiring data (C.S.D., M.F., N.B.P., G.D., A.J.K.), data analysis (C.S.D., N.B.P., A.J.K.), advice regarding scRNAseq data analysis (Y.H., M.S.), writing (C.S.D., M.P.), and reviewing/editing (all authors) the manuscript.

---

**Supplemental Figure 1: Gating strategy for 37 Color T cell cytokine panel. a,** Sample PRI335 (IAI) gated directly *ex vivo*. **b,** Representative gating of phenotypic markers among non-Trm (CD103^-^) without stimulation or **c,** upon stimulation. Populations used for analysis are identified with a color-matched label and box.

**Supplemental Figure 2: Isolating pure iCTB populations from scRNAseq data for downstream analysis. a,** UMAP of single-cell integration of scRNAseq libraries generated from full term placentas at steady state (n=4) and IAI (n=6) following trophoblast enrichment isolation. Trophoblast-containing clusters used for downstream analysis are circled. **b,** Feature plots of trophoblast and immune cell lineage markers; transcripts. **c,** Violin plots depicting expression level of various trophoblast and immune markers. Dashed lines indicate expression cut offs. **d,** UMAP of single-cell integration of cytotrophoblast cells following the removal of cells expressing CD45 and Vimentin >1, rescaling, reharmonizing, and re-clustering for downstream analysis. iCTB-containing clusters used for downstream analysis are circled. **e,** Feature plots of trophoblast and immune cell lineage markers. **f,** UMAP of single-cell integration of cytotrophoblast cells grouped by SS or IAI.

**Supplemental Figure 3: Isolating DPI-specific immune populations from scRNAseq data for downstream analysis. a,** UMAP of single-cell integration of immune cells from full term placentas at steady state (n=6) and IAI (n=6). **b,** UMAP of single-cell integration of immune cells from full term placentas at steady state and IAI separated by tissue type. Clusters containing immune cells of interest including, CD8^+^CD103^+/-^ T cells, FoxP3^+^ T cells, and CD14^+^ immune cells, are circled. **c,** Feature plots of immune cell lineage markers; transcripts and antibody sequencing. **d,** UMAP of single-cell integration of immune cells grouped by SS or IAI.

**Supplemental Figure 4: Isolating DPI-specific T cell populations from scRNAseq data for downstream analysis. a,** UMAP of single-cell integration of immune cells expressing CD8^+^ and CD103^+/-^ protein, following rescaling, reharmonizing, and re-clustering, separated by tissue type. DPI-specific CD8^+^ and CD103^+/-^ clusters used for downstream analysis are circled. **b,** Feature plots of immune cell lineage markers; transcripts and antibody sequencing. **c,** Violin plots depicting expression levels of various immune markers; transcripts and antibody sequencing. Dashed lines indicate expression cut offs. **d,** UMAP of single-cell integration of CD8^+^ and CD103^+/-^ immune cells grouped by steady state or IAI. **e,** UMAP of single-cell integration of immune cells expressing FoxP3 transcript, following rescaling, reharmonizing, and re-clustering, separated by tissue type. DPI-specific FoxP3 clusters used for downstream analysis are circled. **f,** Feature plots of immune cell lineage markers; transcripts and antibody sequencing. **g,** Violin plots depicting expression levels of various immune markers; transcripts and antibody sequencing. **h,** UMAP of single-cell integration of FoxP3+ immune cells grouped by SS or IAI.

**Supplemental Figure 5: Isolating DPI-specific APC populations from scRNAseq data for downstream analysis. a,** UMAP of single-cell integration of immune cells expressing CD14^+^ protein, following rescaling, reharmonizing, and re-clustering, separated by tissue type. DPI-specific CD14^+^ clusters used for downstream analysis are circled. **b,** Feature plots of immune cell lineage markers; transcripts and antibody sequencing. **c,** Violin plots depicting expression levels of various APC immune markers; transcripts and antibody sequencing. **d,** UMAP of single-cell integration of CD14^+^ immune cells grouped by SS or IAI.

**Supplemental Figure 6: Survival and differentiation signaling between T cells and iCTBs is maintained during acute inflammation. a,** Circos plots visualizing the top 25 NicheNet-predicted interactions between T cells and iCTBs during IAI. Interactions with a known role in iCTB survival and differentiation are blue, all others are yellow. Ligands are on the left of each plot, receptors of engaged signaling pathways are on the right. Ligands of specific interest are bold. **b,** Bubble plot depicting average transcript expression level and frequency of predicted T cell ligands. **c,** Circos plots visualizing the top 25 NicheNet-predicted interactions between iCTBs and T cells during IAI. Interactions with a known role in Trm survival and differentiation are blue, all others are yellow. Ligands are on the left of each plot, receptors of engaged signaling pathways are on the right. Ligands of specific interest are bold.

**Supplemental Figure 7: Gating strategy for CTB and APC Panel. a,** Gating strategy for selecting iCTBs and APCs for flow cytometry analysis; cells from single cell trophoblast isolations. **b,** Side scatter (SSC) against individual phenotypic markers and their corresponding positive expression gates. **c,** Black dot = sample PRI325 (SS) used to illustrate gating in a and b. Populations used for analysis are identified with a color-matched label and outline.

**Supplemental Table 1: List of placenta samples and downstream assays.**

## References

1. Janeway CA, Jr. Approaching the asymptote? Evolution and revolution in immunology. Cold Spring Harb Symp Quant Biol. 1989;54 Pt 1:1–13.

2. Chen L, and Flies DB. Molecular mechanisms of T cell co-stimulation and co-inhibition. Nat Rev Immunol. 2013;13(4):227–42.

3. Chen GY, and Nunez G. Sterile inflammation: sensing and reacting to damage. Nat Rev Immunol. 2010;10(12):826–37.

4. Chatterjee S, Do Kang S, Alam S, Salzberg AC, Milici J, van der Burg SH, et al. Tissue-Specific Gene Expression during Productive Human Papillomavirus 16 Infection of Cervical, Foreskin, and Tonsil Epithelium. J Virol. 2019;93(17).

5. Krausgruber T, Fortelny N, Fife-Gernedl V, Senekowitsch M, Schuster LC, Lercher A, et al. Structural cells are key regulators of organ-specific immune responses. Nature. 2020;583(7815):296–302.

6. Nalio Ramos R, Missolo-Koussou Y, Gerber-Ferder Y, Bromley CP, Bugatti M, Nunez NG, et al. Tissue-resident FOLR2(+) macrophages associate with CD8(+) T cell infiltration in human breast cancer. Cell. 2022;185(7):1189–207 e25.

7. Scott MC, Steier Z, Pierson MJ, Stolley JM, O’Flanagan SD, Soerens AG, et al. Deep profiling deconstructs features associated with memory CD8(+) T cell tissue residence. Immunity. 2025;58(1):162–81 e10.

8. Szabo PA. Axes of heterogeneity in human tissue-resident memory T cells. Immunol Rev. 2023;316(1):23–37.

9. Connors TJ, Matsumoto R, Verma S, Szabo PA, Guyer R, Gray J, et al. Site-specific development and progressive maturation of human tissue-resident memory T cells over infancy and childhood. Immunity. 2023;56(8):1894–909.e5.

10. Poon MML, Caron DP, Wang Z, Wells SB, Chen D, Meng W, et al. Tissue adaptation and clonal segregation of human memory T cells in barrier sites. Nat Immunol. 2023;24(2):309–19.

11. Kumar BV, Ma W, Miron M, Granot T, Guyer RS, Carpenter DJ, et al. Human Tissue-Resident Memory T Cells Are Defined by Core Transcriptional and Functional Signatures in Lymphoid and Mucosal Sites. Cell Reports. 2017;20(12):2921–34.

12. Fan X, and Rudensky AY. Hallmarks of Tissue-Resident Lymphocytes. Cell. 2016;164(6):1198–211.

13. Rosato PC, Beura LK, and Masopust D. Tissue resident memory T cells and viral immunity. Curr Opin Virol. 2017;22:44–50.

14. de Visser KE, and Joyce JA. The evolving tumor microenvironment: From cancer initiation to metastatic outgrowth. Cancer Cell. 2023;41(3):374–403.

15. Lui PP, Xu JZ, Aziz H, Sen M, and Ali N. Jagged-1+ skin Tregs modulate cutaneous wound healing. Sci Rep. 2024;14(1):20999.

16. Mathur AN, Zirak B, Boothby IC, Tan M, Cohen JN, Mauro TM, et al. Treg-Cell Control of a CXCL5-IL-17 Inflammatory Axis Promotes Hair-Follicle-Stem-Cell Differentiation During Skin-Barrier Repair. Immunity. 2019;50(3):655–67 e4.

17. Potchen NB, MacMillan HR, Domenjo-Vila E, Konecny AJ, Taber AK, DeJong CS, et al. Tissue-specific adaptation of human T cells is preserved during tissue inflammation. bioRxiv. 2026.

18. Tilburgs T, Claas FHJ, and Scherjon SA. Elsevier Trophoblast Research Award Lecture: Unique Properties of Decidual T Cells and their Role in Immune Regulation during Human Pregnancy. Placenta. 2010;31(SUPPL.):S82–S6.

19. Williams PJ, Searle RF, Robson SC, Innes BA, and Bulmer JN. Decidual leucocyte populations in early to late gestation normal human pregnancy. J Reprod Immunol. 2009;82(1):24–31.

20. Pavličev M, Wagner GP, Chavan AR, Owens K, Maziarz J, Dunn-Fletcher C, et al. Single-cell transcriptomics of the human placenta: Inferring the cell communication network of the maternal-fetal interface. Genome Research. 2017;27(3):349–61.

21. Tsang JCH, Vong JSL, Ji L, Poon LCY, Jiang P, Lui KO, et al. Integrative single-cell and cell-free plasma RNA transcriptomics elucidates placental cellular dynamics. Proceedings of the National Academy of Sciences of the United States of America. 2017;114(37):E7786–E95.

22. Liu Y, Fan X, Wang R, Lu X, Dang YL, Wang H, et al. Single-cell RNA-seq reveals the diversity of trophoblast subtypes and patterns of differentiation in the human placenta. Cell Research. 2018(July).

23. Pique-Regi R, Romero R, Tarca AL, Sendler ED, Xu Y, Garcia-Flores V, et al. Single cell transcriptional signatures of the human placenta in term and preterm parturition. eLife. 2019;8:1–22.

24. Marsh B, Zhou Y, Kapidzic M, Fisher S, and Blelloch R. Regionally distinct trophoblast regulate barrier function and invasion in the human placenta. Elife. 2022;11.

25. Haider S, Lackner AI, Dietrich B, Kunihs V, Haslinger P, Meinhardt G, et al. Transforming growth factor-beta signaling governs the differentiation program of extravillous trophoblasts in the developing human placenta. Proc Natl Acad Sci U S A. 2022;119(28):e2120667119.

26. Li Q, Sharkey A, Sheridan M, Magistrati E, Arutyunyan A, Huhn O, et al. Human uterine natural killer cells regulate differentiation of extravillous trophoblast early in pregnancy. Cell Stem Cell. 2024;31(2):181–95 e9.

27. Monsivais D, Clementi C, Peng J, Fullerton PT, Jr., Prunskaite-Hyyrylainen R, Vainio SJ, et al. BMP7 Induces Uterine Receptivity and Blastocyst Attachment. Endocrinology. 2017;158(4):979–92.

28. Nagashima T, Li Q, Clementi C, Lydon JP, DeMayo FJ, and Matzuk MM. BMPR2 is required for postimplantation uterine function and pregnancy maintenance. J Clin Invest. 2013;123(6):2539–50.

29. Peters Q, and Prlic M. Six degrees of TGFbeta. Nat Immunol. 2025;26(8):1221–2.

30. Devi KSP, Wang E, Jaiswal A, Konieczny P, Kim TG, Nirschl CJ, et al. PD-1 is requisite for skin T(RM) cell formation and specification by TGFbeta. Nat Immunol. 2025;26(8):1339–51.

31. Jain VG, Willis KA, Jobe A, and Ambalavanan N. Chorioamnionitis and neonatal outcomes. Pediatr Res. 2022;91(2):289–96.

32. Grebenciucova E, and VanHaerents S. Interleukin 6: at the interface of human health and disease. Front Immunol. 2023;14:1255533.

33. Jacobsson B, Mattsby-Baltzer I, and Hagberg H. Interleukin-6 and interleukin-8 in cervical and amniotic fluid: relationship to microbial invasion of the chorioamniotic membranes. BJOG. 2005;112(6):719–24.

34. Martin KR, Wong HL, Witko-Sarsat V, and Wicks IP. G-CSF - A double edge sword in neutrophil mediated immunity. Semin Immunol. 2021;54:101516.

35. Raeber ME, Zurbuchen Y, Impellizzieri D, and Boyman O. The role of cytokines in T-cell memory in health and disease. Immunol Rev. 2018;283(1):176–93.

36. Monsivais D, Nagashima T, Prunskaite-Hyyrylainen R, Nozawa K, Shimada K, Tang S, et al. Endometrial receptivity and implantation require uterine BMP signaling through an ACVR2A-SMAD1/SMAD5 axis. Nat Commun. 2021;12(1):3386.

37. Bai X, Williams JL, Greenwood SL, Baker PN, Aplin JD, and Crocker IP. A placental protective role for trophoblast-derived TNF-related apoptosis-inducing ligand (TRAIL). Placenta. 2009;30(10):855–60.

38. Straszewski-Chavez SL, Abrahams VM, and Mor G. The role of apoptosis in the regulation of trophoblast survival and differentiation during pregnancy. Endocr Rev. 2005;26(7):877–97.

39. Fock V, Plessl K, Draxler P, Otti GR, Fiala C, Knofler M, et al. Neuregulin-1-mediated ErbB2-ErbB3 signalling protects human trophoblasts against apoptosis to preserve differentiation. J Cell Sci. 2015;128(23):4306–16.

40. Chen X, Qi L, Zhao C, Xue J, Chen M, Diao L, et al. Decreased expression of SEMA4D in recurrent implantation failure induces reduction of trophoblast invasion and migration via the Met/PI3K/Akt pathway. J Reprod Immunol. 2022;153:103657.

41. Li G, Ma L, Lu H, Cao G, Shao X, Liu Y, et al. Transactivation of Met signalling by semaphorin4D in human placenta: implications for the pathogenesis of preeclampsia. J Hypertens. 2018;36(11):2215–25.

42. Boraschi D, Italiani P, Weil S, and Martin MU. The family of the interleukin-1 receptors. Immunol Rev. 2018;281(1):197–232.

43. Banerjee S, Smallwood A, Moorhead J, Chambers AE, Papageorghiou A, Campbell S, et al. Placental expression of interferon-gamma (IFN-gamma) and its receptor IFN-gamma R2 fail to switch from early hypoxic to late normotensive development in preeclampsia. J Clin Endocrinol Metab. 2005;90(2):944–52.

44. Rizzuto G, Brooks JF, Tuomivaara ST, McIntyre TI, Ma S, Rideaux D, et al. Establishment of fetomaternal tolerance through glycan-mediated B cell suppression. Nature. 2022;603(7901):497–502.

45. Fan X, Muruganandan S, Shallie PD, Dhal S, Petitt M, and Nayak NR. VEGF Maintains Maternal Vascular Space Homeostasis in the Mouse Placenta through Modulation of Trophoblast Giant Cell Functions. Biomolecules. 2021;11(7).

46. Fan X, Rai A, Kambham N, Sung JF, Singh N, Petitt M, et al. Endometrial VEGF induces placental sFLT1 and leads to pregnancy complications. J Clin Invest. 2014;124(11):4941–52.

47. Zhou Y, McMaster M, Woo K, Janatpour M, Perry J, Karpanen T, et al. Vascular endothelial growth factor ligands and receptors that regulate human cytotrophoblast survival are dysregulated in severe preeclampsia and hemolysis, elevated liver enzymes, and low platelets syndrome. Am J Pathol. 2002;160(4):1405–23.

48. Li W, Li H, Bocking AD, and Challis JR. Tumor necrosis factor stimulates matrix metalloproteinase 9 secretion from cultured human chorionic trophoblast cells through TNF receptor 1 signaling to IKBKB-NFKB and MAPK1/3 pathway. Biol Reprod. 2010;83(3):481–7.

49. Hannan NJ, Jones RL, White CA, and Salamonsen LA. The chemokines, CX3CL1, CCL14, and CCL4, promote human trophoblast migration at the feto-maternal interface. Biol Reprod. 2006;74(5):896–904.

50. Galeano Nino JL, Pageon SV, Tay SS, Colakoglu F, Kempe D, Hywood J, et al. Cytotoxic T cells swarm by homotypic chemokine signalling. Elife. 2020;9.

51. Zhang S, Ding J, Zhang Y, Liu S, Yang J, and Yin T. Regulation and Function of Chemokines at the Maternal-Fetal Interface. Front Cell Dev Biol. 2022;10:826053.

52. Huang Y, Zhu XY, Du MR, and Li DJ. Human trophoblasts recruited T lymphocytes and monocytes into decidua by secretion of chemokine CXCL16 and interaction with CXCR6 in the first-trimester pregnancy. J Immunol. 2008;180(4):2367–75.

53. Yeo J, Ko M, Lee DH, Park Y, and Jin HS. TIGIT/CD226 Axis Regulates Anti-Tumor Immunity. Pharmaceuticals (Basel). 2021;14(3).

54. Yoshida H, and Hunter CA. The immunobiology of interleukin-27. Annu Rev Immunol. 2015;33:417–43.

55. Wang G, Munoz-Rojas AR, Spallanzani RG, Franklin RA, Benoist C, and Mathis D. Adipose-tissue Treg cells restrain differentiation of stromal adipocyte precursors to promote insulin sensitivity and metabolic homeostasis. Immunity. 2024;57(6):1345–59 e5.

56. Ali N, and Rosenblum MD. Regulatory T cells in skin. Immunology. 2017;152(3):372–81.

57. Maryanovich M, and Frenette PS. T-Regulating Hair Follicle Stem Cells. Immunity. 2017;46(6):979–81.

58. Ali N, Zirak B, Rodriguez RS, Pauli ML, Truong HA, Lai K, et al. Regulatory T Cells in Skin Facilitate Epithelial Stem Cell Differentiation. Cell. 2017;169(6):1119–29 e11.

59. Munoz-Rojas AR, and Mathis D. Tissue regulatory T cells: regulatory chameleons. Nat Rev Immunol. 2021;21(9):597–611.

60. Luan J, Truong C, Vuchkovska A, Guo W, Good J, Liu B, et al. CD80 on skin stem cells promotes local expansion of regulatory T cells upon injury to orchestrate repair within an inflammatory environment. Immunity. 2024;57(5):1071–86 e7.

61. Gebhardt T, Wakim LM, Eidsmo L, Reading PC, Heath WR, and Carbone FR. Memory T cells in nonlymphoid tissue that provide enhanced local immunity during infection with herpes simplex virus. Nat Immunol. 2009;10(5):524–30.

62. Jiang X, Clark RA, Liu L, Wagers AJ, Fuhlbrigge RC, and Kupper TS. Skin infection generates non-migratory memory CD8+ TRM cells providing global skin immunity. Nature. 2012;483(7388):227–31.

63. Trundley A, and Moffett A. Human uterine leukocytes and pregnancy. Tissue Antigens. 2004;63(1):1–12.

64. Hanna J, Goldman-Wohl D, Hamani Y, Avraham I, Greenfield C, Natanson-Yaron S, et al. Decidual NK cells regulate key developmental processes at the human fetal-maternal interface. Nat Med. 2006;12(9):1065–74.

65. Moffett-King A. Natural killer cells and pregnancy. Nat Rev Immunol. 2002;2(9):656–63.

66. Mackay LK, Rahimpour A, Ma JZ, Collins N, Stock AT, Hafon ML, et al. The developmental pathway for CD103(+)CD8+ tissue-resident memory T cells of skin. Nat Immunol. 2013;14(12):1294–301.

67. Mohammed J, Beura LK, Bobr A, Astry B, Chicoine B, Kashem SW, et al. Stromal cells control the epithelial residence of DCs and memory T cells by regulated activation of TGF-β. Nature Immunology. 2016;17(4):414–21.

68. Mani V, Bromley SK, Äijö T, Mora-Buch R, Carrizosa E, Warner RD, et al. Migratory DCs activate TGF-b to precondition naïve CD8+T cells for tissue-resident memory fate. Science. 2019;366(6462).

69. Kim J, Chang DY, Lee HW, Lee H, Kim JH, Sung PS, et al. Innate-like Cytotoxic Function of Bystander-Activated CD8 + T Cells Is Associated with Liver Injury in Acute Hepatitis A. Immunity. 2018;48(1):161–73.e5.

70. Liu Y, Beyer A, and Aebersold R. On the Dependency of Cellular Protein Levels on mRNA Abundance. Cell. 2016;165(3):535–50.

71. Maurice NJ, Erickson JR, DeJong CS, Mair F, Taber AK, Frutoso M, et al. Cytokine and metabolite networks shape T cell residency and functionality at the term human maternal-fetal interface. J Immunol. 2025.

72. Hunkapiller NM, and Fisher SJ. Chapter 12. Placental remodeling of the uterine vasculature. Methods Enzymol. 2008;445:281–302.

73. Erickson JR, Mair F, Bugos G, Martin J, Tyznik AJ, Nakamoto M, et al. AbSeq Protocol Using the Nano-Well Cartridge-Based Rhapsody Platform to Generate Protein and Transcript Expression Data on the Single-Cell Level. STAR Protoc. 2020;1(2).

74. Hao Y, Hao S, Andersen-Nissen E, Mauck WM, 3rd, Zheng S, Butler A, et al. Integrated analysis of multimodal single-cell data. Cell. 2021;184(13):3573–87 e29.

75. Browaeys R, Saelens W, and Saeys Y. NicheNet: modeling intercellular communication by linking ligands to target genes. Nat Methods. 2020;17(2):159–62.

76. Liechti T, Weber LM, Ashhurst TM, Stanley N, Prlic M, Van Gassen S, et al. An updated guide for the perplexed: cytometry in the high-dimensional era. Nat Immunol. 2021;22(10):1190–7.

77. Mair F, and Tyznik AJ. High-Dimensional Immunophenotyping with Fluorescence-Based Cytometry: A Practical Guidebook. Methods Mol Biol. 2019;2032:1–29.

78. Mair F, and Prlic M. OMIP-044: 28-color immunophenotyping of the human dendritic cell compartment. Cytometry A. 2018;93(4):402–5.

79. Frutoso M, DeJong CS, Shree R, McCartney SA, and Prlic M. Phenotypically similar but functionally distinct NK cell populations within the human maternal-fetal interface. Immunohorizons. 2026;10(3).

