## Supplemental Figures for "Human tissue-resident CD8 T cells contribute to trophoblast homeostasis in health and during acute inflammation"

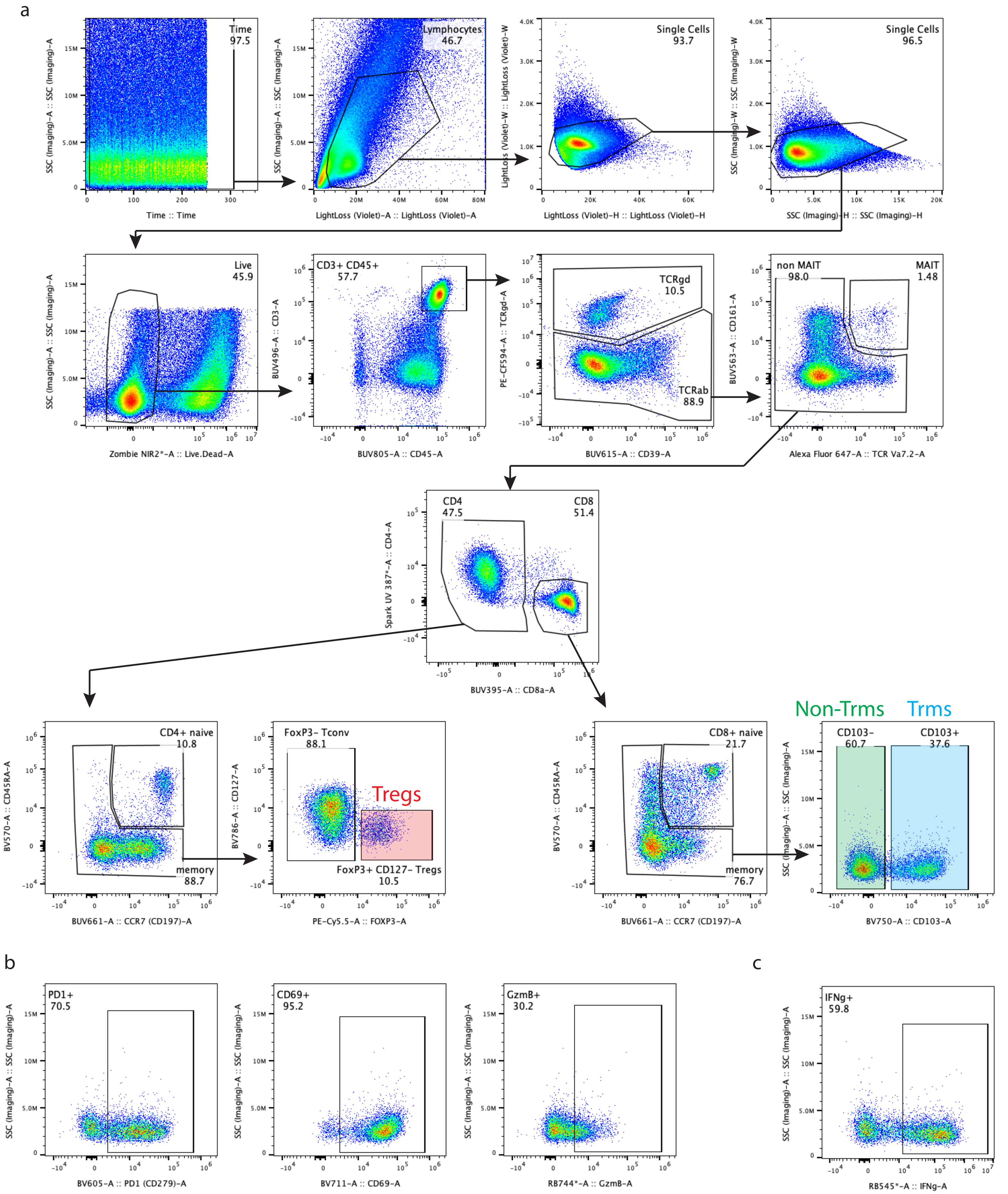


**Supplemental Figure 1: Gating strategy for 37 Color T cell cytokine panel.** a, Sample mfPLAC335 (IAI) gated directly ex vivo. b, Representative gating of phenotypic markers among non-TRMs (CD103-) without stimulation or c, upon stimulation. Populations used for analysis are identified with a color-matched label and box.

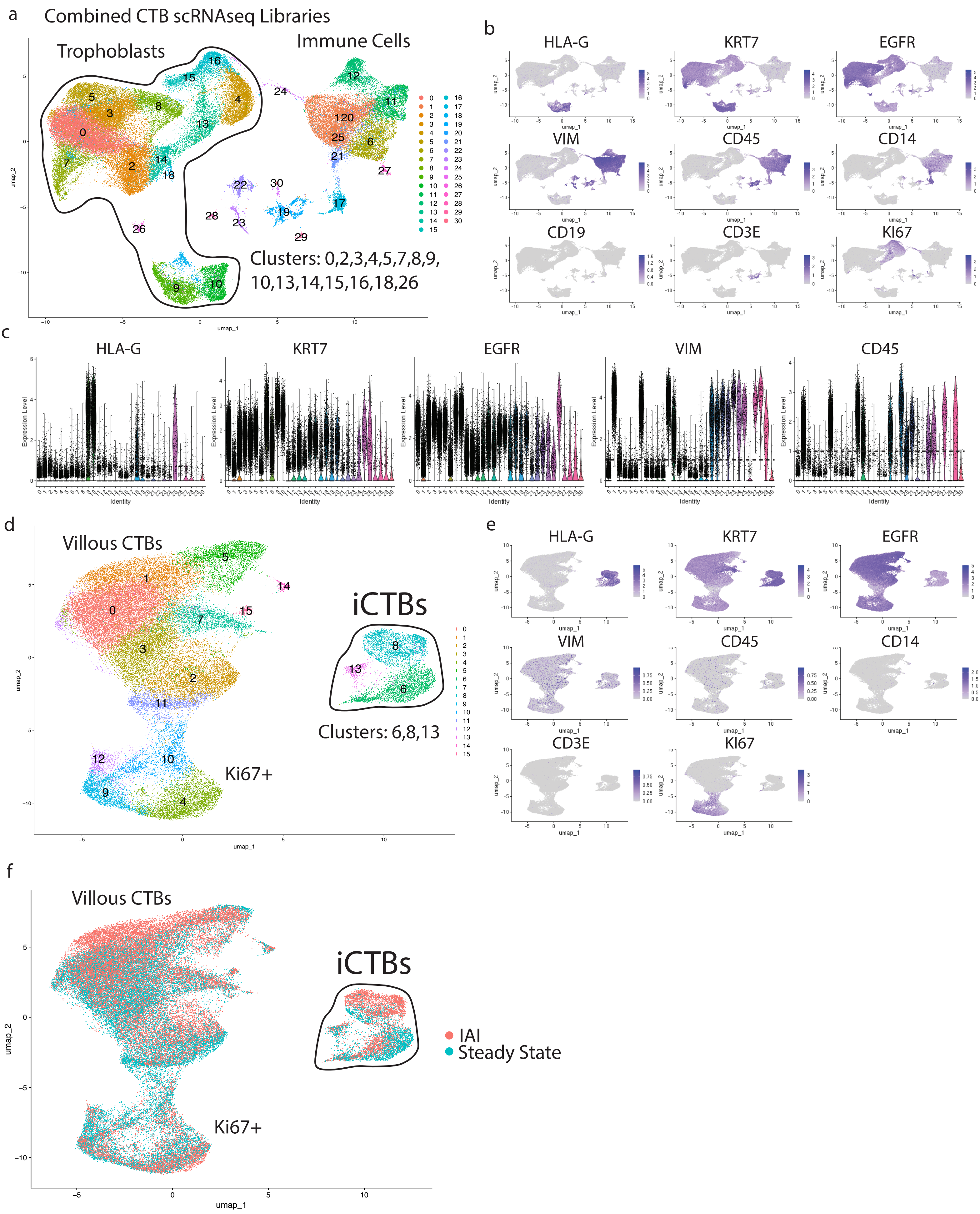


**Supplemental Figure 2: Isolating pure iCTB populations from scRNAseq data for downstream analysis.**

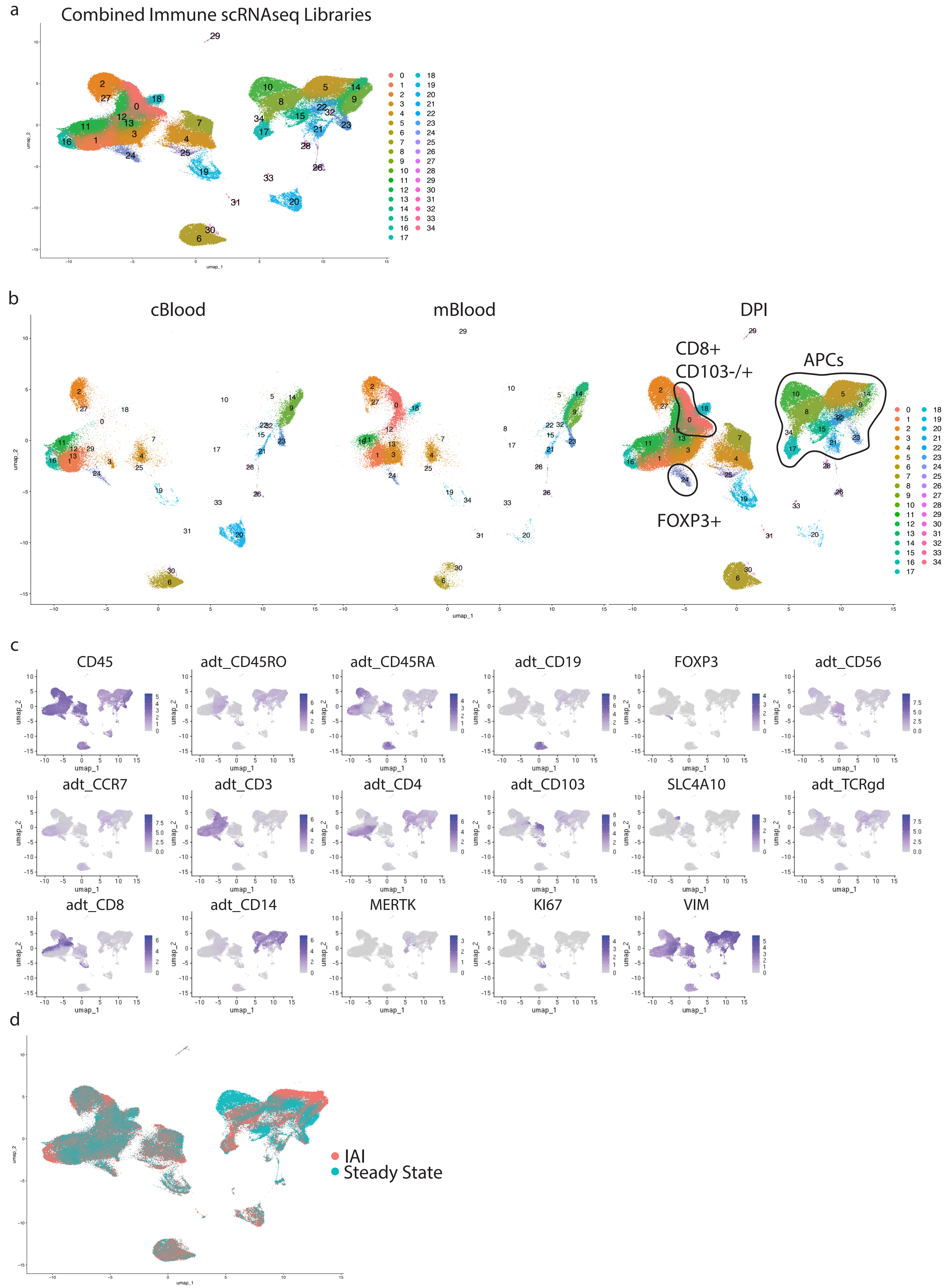


**Supplemental Figure 3: Isolating DPI-specific immune populations from scRNAseq data for downstream analysis.**

### DPI-specific Trm and Non-Trm CD8+ T cells

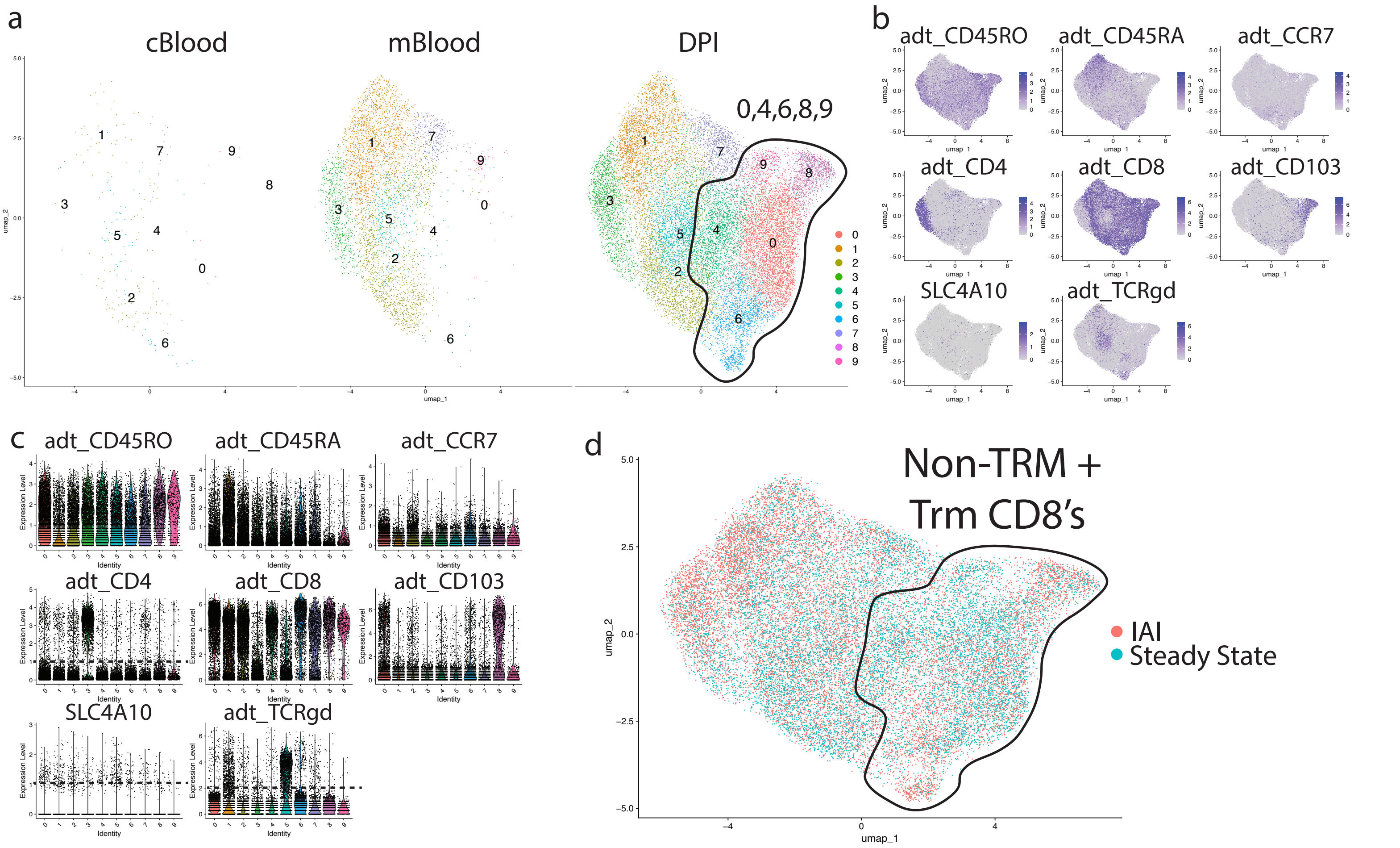


### PDI-specific Tregs

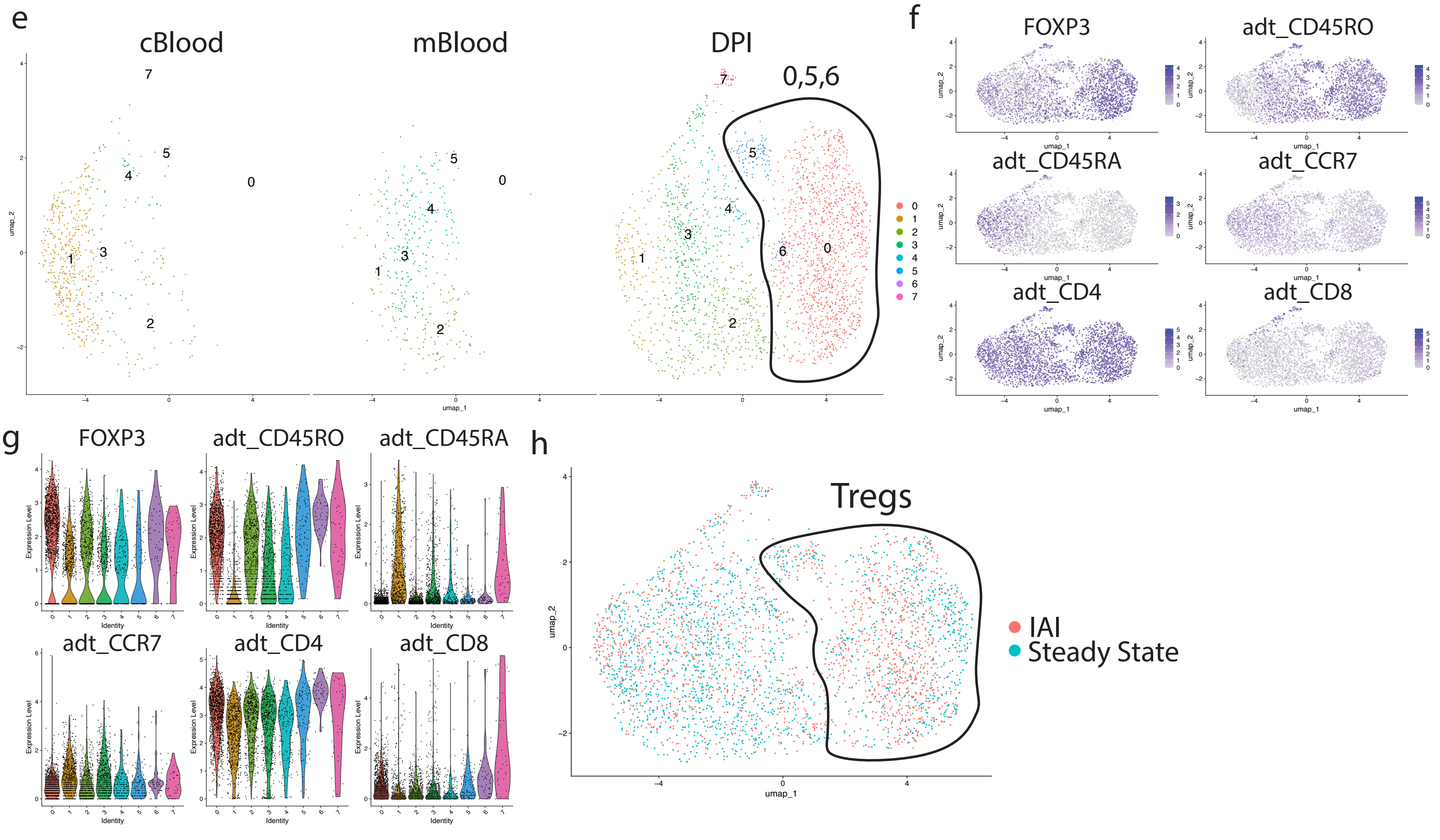


**Supplemental Figure 4: Isolating DPI-specific T cell populations from scRNAseq data for downstream analysis.**

### DPI-specific APCs

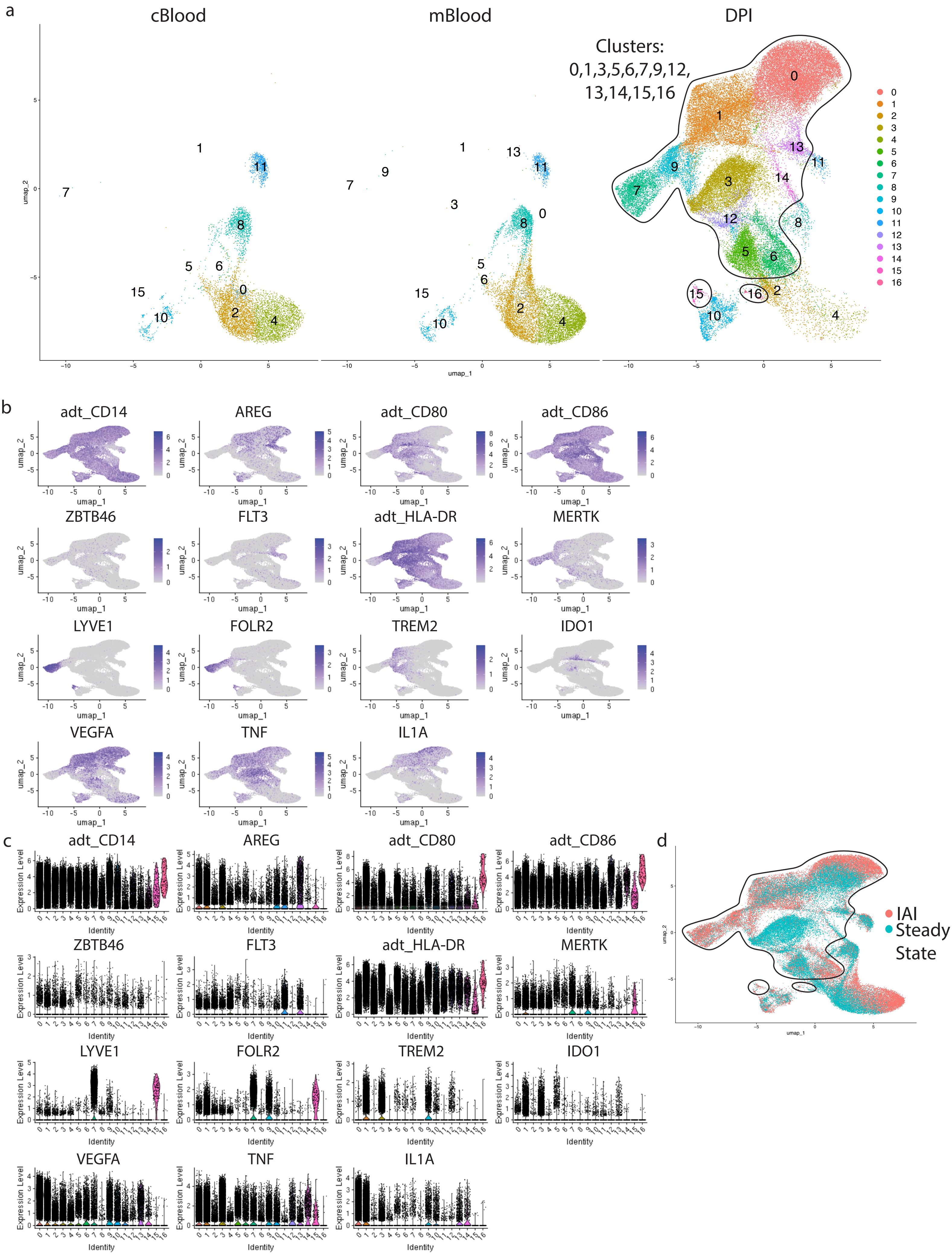


**Supplemental Figure 5: Isolating DPI-specific APC populations from scRNAseq data for downstream analysis.**

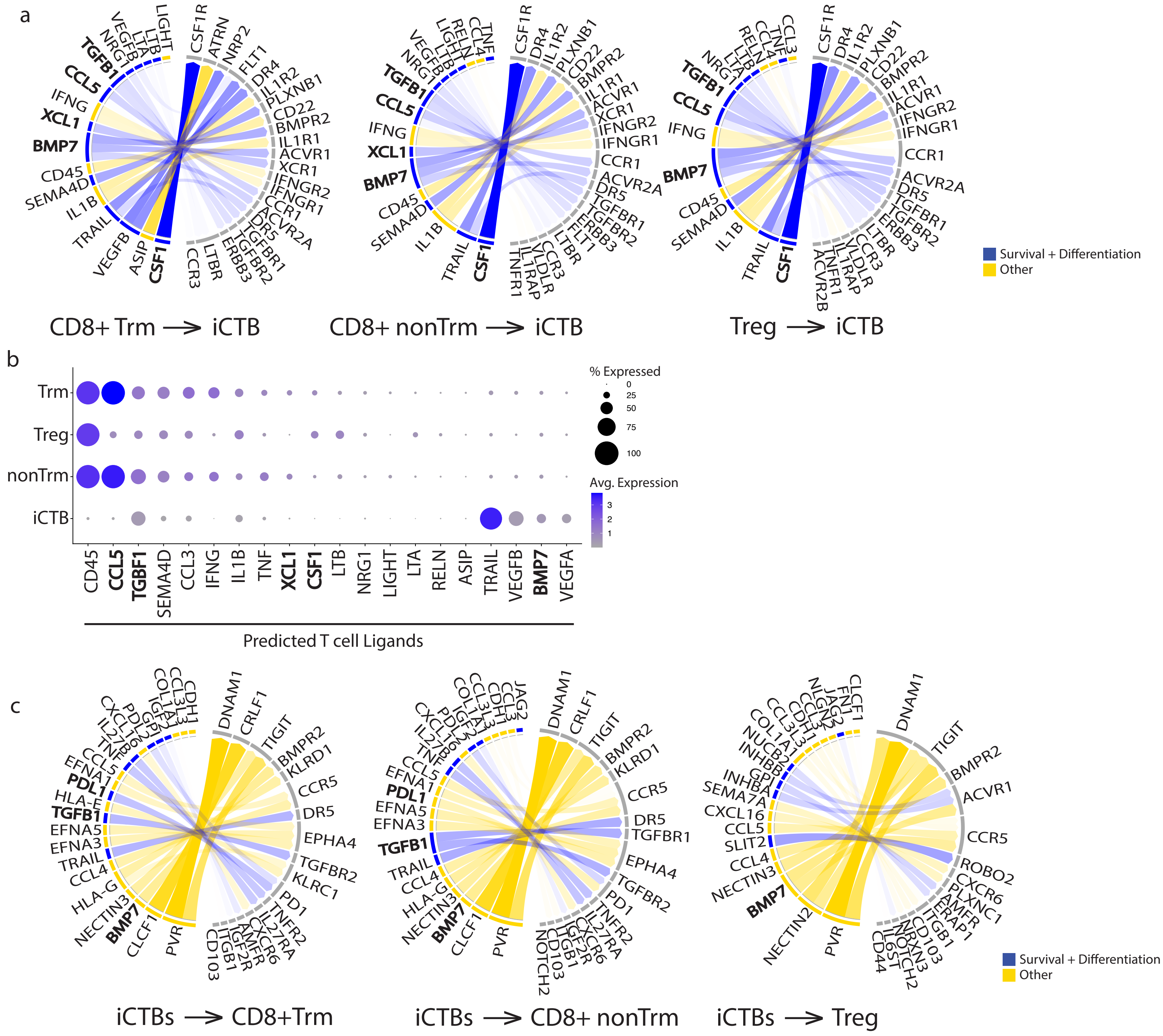


**Supplemental Figure 6: Survival and differentiation signaling between T cells and iCTB is maintained during acute inflammation.**

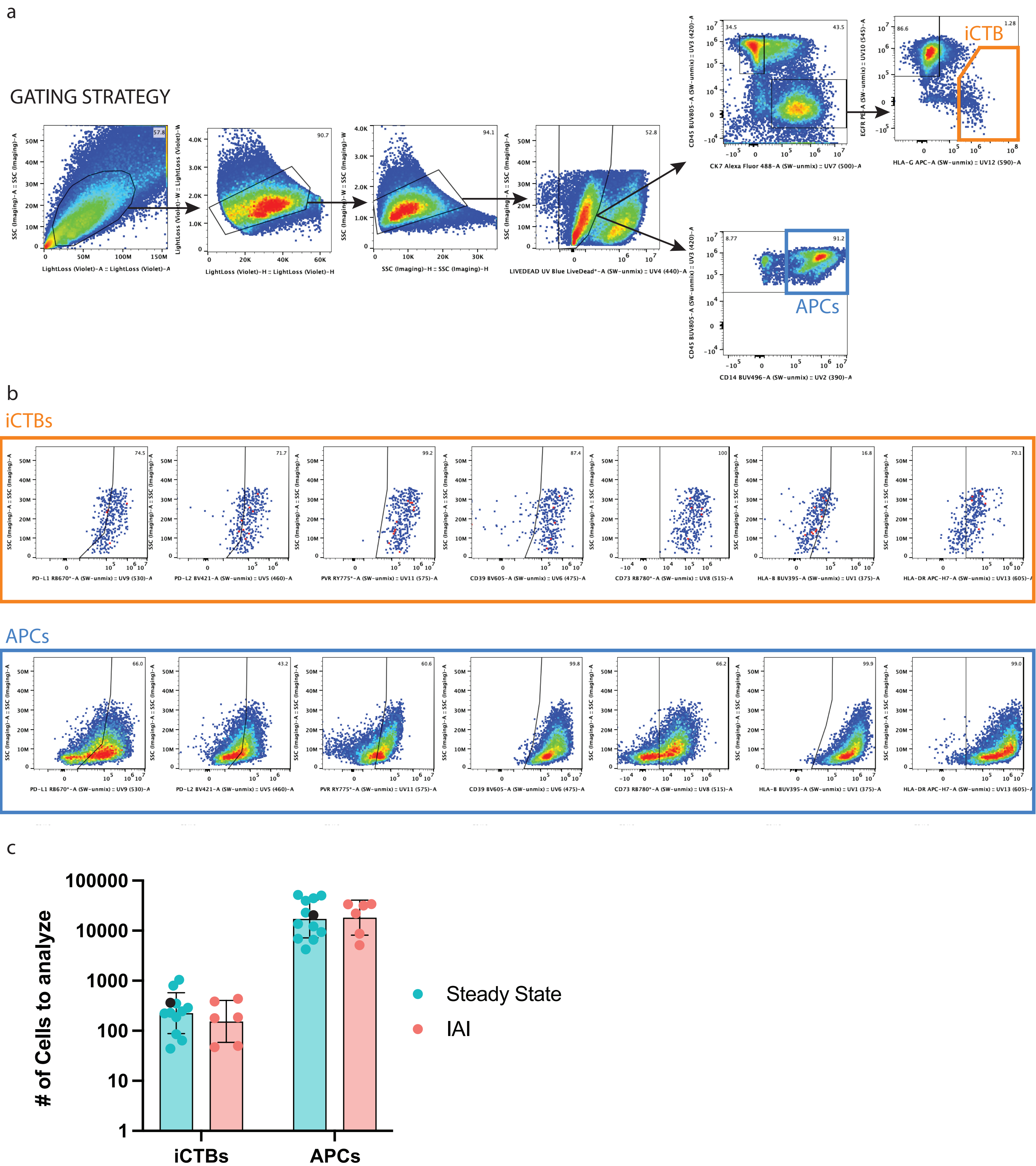


**Supplemental Figure 7: Gating strategy for CTB and APC Panel.**
