## Supplemental Table 1 for "Human tissue-resident CD8 T cells contribute to trophoblast homeostasis in health and during acute inflammation"

**List of placental samples**

| Study ID# | Age | | GA Delivery | Delivery | Labor | Fetal Sex | IAI |
|---|---|---|---|---|---|---|---|
| PRI271 | 29 | | 39w3d | CS | No | F | No |
| PRI282 | 33 | | 41w6d | CS | Yes | M | No |
| PRI284 | 29 | | 40w0d | VD | Yes | F | Yes |
| PRI286 | 28 | | 39w2d | CS | No | F | No |
| PRI289 | 35 | | 39w5d | CS | No | F | No |
| PRI292 | 18 | | 39w5d | VD | Yes | F | Yes |
| PRI300 | 34 | | 39w1d | CS | No | F | No |
| PRI302 | 35 | | 40w2d | CS | Yes | M | Yes |
| PRI303 | 38 | | 36w3d | CS | No | F | No |
| PRI304 | 35 | | 39w3d | CS | No | M | No |
| PRI305 | 33 | | 40w4d | CS | Yes | F | Yes |
| PRI306 | 34 | | 39w5d | CS | Yes | F | Yes |
| PRI307 | 32 | | 39w3d | CS | No | F | No |
| PRI308 | 38 | | 39w0d | VD | Yes | F | Yes |
| PRI309 | 36 | | 40w0d | VD | Yes | M | Yes |
| PRI310 | 31 | | 40w0d | CS | Yes | M | Yes |
| PRI313 | 32 | | 37w1d | CS | No | F | No |
| PRI315 | 35 | | 39w2d | CS | No | M | No |
| PRI316 | 39 | | 39w1d | CS | No | | No |
| PRI317 | 43 | | 37w0d | CS | No | M | No |
| PRI318 | 46 | | 35w0d | CS | No | M | No |
| PRI319 | 37 | | 37w4d | CS | No | F | No |
| PRI320 | 30 | | 37w1d | CS | No | | No |
| PRI321 | 34 | | 40w2d | VD | Yes | F | Yes |
| PRI325 | 36 | | 39w4d | CS | No | F | No |
| PRI331 | 37 | | 39w4d | CS | Yes | M | Yes |
| PRI333 | 26 | | 37w6d | VD | Yes | M | No |
| PRI335 | 38 | | 40w0d | CS | Yes | M | Yes |
| PRI338 | 32 | | 39w1d | VD | Yes | M | Yes |
| PRI339 | 35 | | 39w4d | CS | No | M | No |
| PRI340 | 33 | | 39w0d | CS | Yes | M | Yes |
| PRI342 | 35 | | 41w2d | VD | Yes | F | No |
| PRI344 | 36 | | 39w4d | VD | Yes | M | No |
| PRI351 | 34 | | 39w2d | CS | No | M | No |
| PRI354 | 25 | | 39w6d | VD | Yes | F | Yes |

G/P = Gravita/Parity ; GA = gestational age ; IAI = Intra-amniotic inflammation ; CS = cesarian section ; VD = vaginal delivery
